# De novo cholesterol biosynthesis in the bacterial domain

**DOI:** 10.1101/2022.10.03.510737

**Authors:** Alysha K. Lee, Jeremy H. Wei, Paula V. Welander

**Affiliations:** Department of Earth System Science, Stanford University, Stanford, CA 94305

## Abstract

Sterols are versatile lipids primarily associated with eukaryotes. While bacteria also produce sterols and studies of bacterial biosynthesis proteins have revealed novel biosynthetic pathways and a potential evolutionary role in the origin of sterol biosynthesis, no bacterium has been shown to synthesize highly modified eukaryotic sterols, such as cholesterol. This has led to the notion that bacteria only produce biosynthetically simple sterols and has lessened the consideration of bacterial production in discussions of sterol biosynthesis. In this study, we demonstrate two phylogenetically distinct bacteria, *Enhygromyxa salina* and *Calothrix* sp. NIES-4105, are capable of *de novo* cholesterol production. We also identified 25-hydroxycholesterol released as a product of acid hydrolysis in extracts from both bacteria, suggesting cholesterol exists as a conjugated molecule in these organisms. We coupled our lipid extractions to bioinformatic analyses and heterologous expression experiments to identify genetic pathways driving cholesterol production in each bacterium. *E. salina* shares much of its cholesterol biosynthesis pathway with the canonical eukaryotic pathway, except for C-4 demethylation where we identified a unique variation on the bacterial C-4 demethylation pathway. *Calothrix* lacks homologs for several steps in cholesterol biosynthesis, suggesting this bacterium may harbor a novel mechanism for completing cholesterol biosynthesis. Altogether, these results demonstrate the complexity underpinning bacterial sterol biosynthesis and raise further questions about the functional and regulatory roles of sterols in bacteria.

## Introduction

Sterols are a class of ubiquitous and essential eukaryotic lipids important in a variety of physiological functions including cell signaling, membrane homeostasis, and developmental timing (1)(2)(3)(4). Eukaryotic sterol biosynthesis has been studied extensively revealing complex biosynthetic pathways including a shared set of enzymes used to produce similar sterol end-products that vary in the level of unsaturations, demethylations, and alkalytions (5)(6). The order of reactions in these biosynthetic pathways dictate the production and accumulation of sterol intermediates which play a regulatory role in lipid homeostasis (7)(8), downstream product biosynthesis (9), and stress response (10) (11). The chemical modifications required to synthesize cholesterol in vertebrates, phytosterols in plants, and ergosterol in fungi are essential to the biophysical properties of these lipids, effecting localization and membrane dynamics in their respective organisms (12) (13) (14). These sterols also serve as a branching point for downstream metabolite biosynthesis. Oxidation reactions are involved in converting sterols into a wide range of compounds including oxysterols, bile acids, steroid hormones, and brassinosteroids which can all function as ligands in various signaling pathways (15)(16)(17). Eukaryotes also conjugate sterols to sugars, proteins, or other lipids, further expanding the function of these lipids to include cell defense, energy storage, and signaling (18)(19)(20). Overall, the complexity of eukaryotic sterol biosynthesis is reflective of the diverse functions and significant roles these lipids play in eukaryotic physiology.

While sterol biosynthesis and function has been well-studied in eukaryotic organisms, bacterial sterol synthesis and function is comparatively underexplored. Several bacteria, including aerobic methanotrophs, Planctomycetes, and various myxobacteria are known to produce sterols *de novo* (21). Unlike eukaryotes, these bacteria largely produce lanosterol, parkeol, or cycloartenol, the initial cyclization products of oxidosqualene cyclase (OSC)(22)(23)(24). However, some bacteria perform additional chemical modifications during sterol synthesis; methanotrophic bacteria modify sterols into distinct monomethylated structures specific to Methylococcaceae (21), sterol-producing Planctomycetes conjugate sterols to an unidentified macromolecule (23), and several Myxococcota (25) produce intermediates in the cholesterol biosynthesis pathway, including zymosterol (21)(26)(24). Furthermore, phylogenomic studies have expanded the number of potential bacterial sterol producers, identifying the genes required for both cyclization and downstream modifications in phyla across the bacterial domain (27) (28). Several of these bacteria harbor the genetic potential to produce the biosynthetically complex sterols associated with eukaryotes, including cholesterol, however lipid analyses have yet to detect these sterols in bacteria (6).

The discrepancy between the genomic capacity for complex sterol production and the observed sterols in bacteria prompted us to undertake more comprehensive lipid analyses of sterol-producing bacteria. We hypothesized previous analyses may have underestimated the bacterial sterol inventory due to limited biomass and/or inadequate lipid extraction techniques. Here, we demonstrate the marine myxobacterium *Enhygromyxa salina* and the cyanobacterium *Calothrix* sp. NIES-4105 are both capable of producing cholesterol. We identified homologs to some, but not all, of the steps in eukaryotic cholesterol biosynthesis, differentiating bacterial cholesterol production from eukaryotes. We also demonstrate that demethylation at C-4, a crucial step in cholesterol biosynthesis, occurs by a distinct mechanism in each bacterium. Our analyses of sterol biosynthesis in these phylogenetically distinct bacteria suggest a complex biology underpinning bacterial cholesterol production and raise broader questions about sterol biosynthesis, evolution, and function.

## Results

### *Enhygromyxa salina* produces free and conjugated sterols

We previously demonstrated the production of zymosterol, a biosynthetic intermediate in cholesterol synthesis, in lipid extracts of the myxobacterium *Enhygromyxa salina* (21). However, *E. salina* is a social predatory bacterium cultured on solid agar, limiting the amount of biomass available for extensive lipid analyses (29). These culturing conditions also present a potential source of sterol contamination as both agar and supplemental yeast may contain sterols. To better assess the sterol inventory of this organism, we adapted *E. salina* to growth in a liquid medium supplemented with whole cell *Escherichia coli*, which does not natively produce sterols. These culturing conditions increased extractable lipids 50-fold, allowing for more extensive analyses.

We first extracted free lipids from *E. salina* biomass through a Bligh-Dyer extraction (30). These initial analyses revealed a variety of sterols, including cholesterol (Fig. 1a; *SI Appendix*, Fig. S1). In addition to cholesterol, we detected 4,4-dimethyl(zymo)sterol, an intermediate only demethylated at C-14, indicating *E. salina* performs C-14 demethylation before C-4 demethylation, like fungi and animals, and not after removal of the first C-4 methyl group, like plants (6). We also detected several downstream intermediates unsaturated at C-24 including zymosterol, dehydrolathosterol, and desmosterol but no intermediates saturated at C-24. The accumulation of only C-24 unsaturated intermediates suggests *E. salina* favors the Bloch cholesterol biosynthesis pathway, where C-24 reduction occurs as the final step in cholesterol biosynthesis (*SI Appendix*, Fig. S2) (31). Finally, to control for potential contamination, we extracted both culturing medium and concentrated supplemental *E. coli* without bacterial biomass. Both the media and extraction controls were devoid of sterols (*SI Appendix*, Fig. S3).

**Fig. 1.**
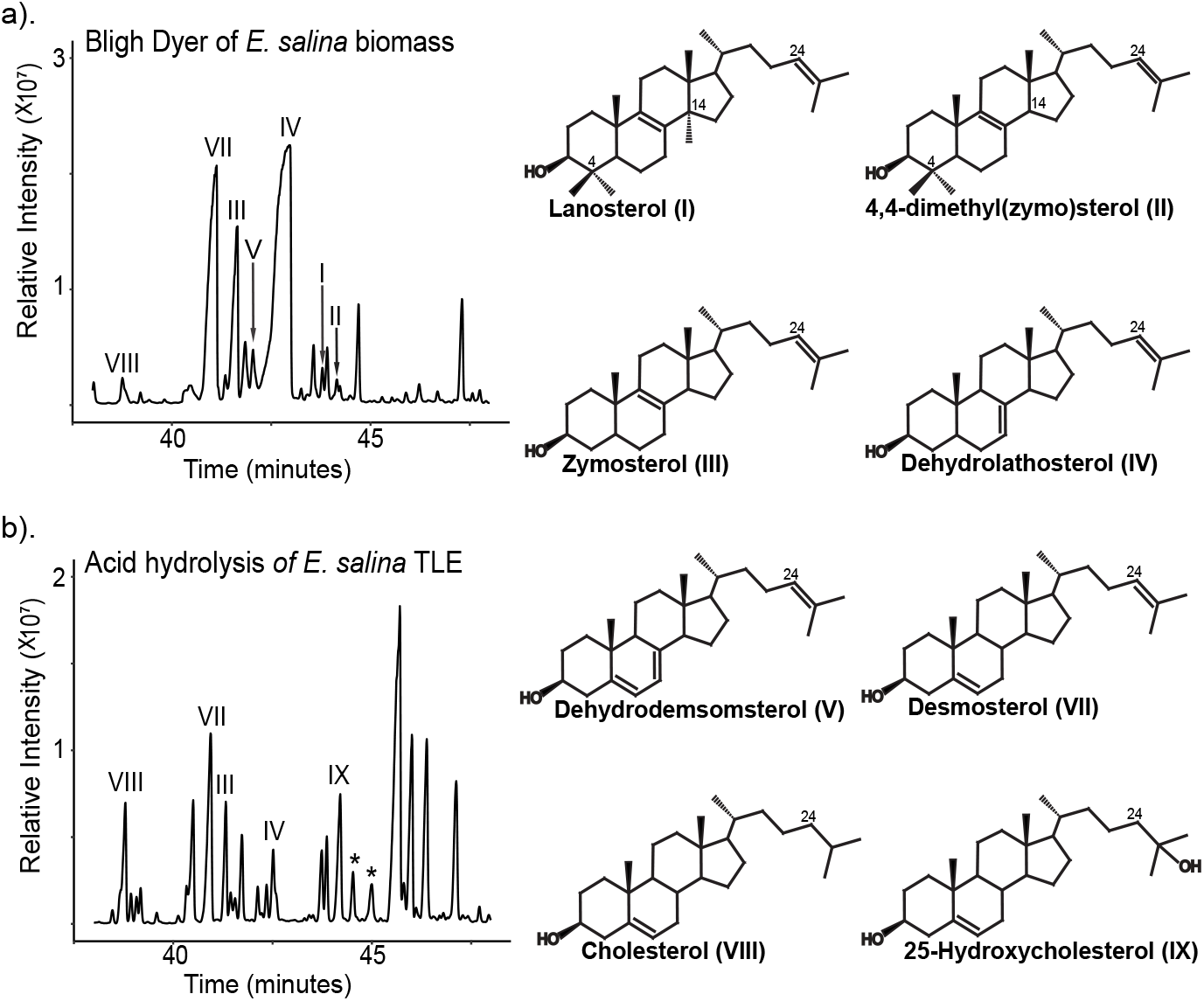
*E. salina* synthesizes unbound and bound sterols. (a) Total ion chromatogram of free sterols from *E. salina* extracted using modified Bligh Dyer procedure. Identified sterols included cholesterol (VIII) as well as C-24 unsaturated intermediates (I-VII). (b) Total ion chromatogram of ether and ester bound sterols released from *E. salina* lipid extracts by acid hydrolysis. Hydrolysis released additional sterols, including 25-hydroxycholesterol (IX) and other putative hydroxysterols (*). All lipids were derivatized to trimethylsilyl groups. Mass spectra of identified sterols are shown in *SI Appendix*, Fig S1.

To assess *E. salina* for potential sterol conjugates, we hydrolyzed both total lipid extract (TLE) and cell biomass with either methanolic base, which cleaves ester bound lipids, or methanolic acid, which cleaves ester and ether bound lipids. Hydrolysis of TLE with methanolic base did not release additional sterols but hydrolysis with methanolic acid released 25-hydroxycholesterol (25-OHC) and two other probable oxysterols (Fig. 1b; *SI Appendix*, Fig. S4). The presence of oxysterols after acid hydrolysis of TLE, but not after base hydrolysis, suggests that these conjugations are mediated by an ether bond. To further characterize the chemical nature of sterol conjugates, we separated extractable lipids by polarity using Si-gel column chromatography and hydrolyzed fractions with methanolic acid. After hydrolysis, all sterols, including hydroxysterols, were only present in the alcohol fraction, suggesting conjugates are similar in polarity to free sterols. Direct hydrolysis of cell biomass with either methanolic base or acid did not significantly alter the *E. salina* sterol profile from that of the hydrolyzed TLEs. To confirm these additional sterols were not the products of degradation or autooxidation during hydrolysis, we acid hydrolyzed cholesterol and desmosterol standards and did not detect production of any hydroxysterols (*SI Appendix*, Fig. S5).

### *E. salina* harbors eukaryotic cholesterol biosynthesis proteins

Production of cholesterol by *E. salina* prompted us to search for a putative biosynthesis pathway in its genome. We conducted a BLASTp search (<1 x e^-30^, 30% ID) against sterol biosynthesis proteins from eukaryotes and bacteria (5)(32). We chose a restrictive cut-off for our BLASTp search to ensure identified proteins are involved in sterol biosynthesis, as proteins in the cholesterol biosynthesis pathway belong to large superfamilies that include functions outside sterol biosynthesis (6). We detected homologs for nearly every step in the eukaryotic cholesterol biosynthesis pathway in the *E. salina* genome (Fig 2a; *SI Appendix*, Table S1). Notably, sterol biosynthesis genes are not localized in gene clusters, as observed in other sterol-producing bacteria (33), but instead found in separate loci across the genome (*SI Appendix*, Fig. S6). Using this newly constructed cholesterol biosynthesis pathway, we next searched for homologs in other bacteria with demonstrated genomic capacity for sterol production (<1 x e^-50^, 30% ID). We identified 103 bacterial isolates and 21 metagenome assembled genomes with squalene monooxygenase (SMO) and oxidosqualene cyclase (OSC), the first committed steps in sterol biosynthesis. Of these bacteria, only other sequenced isolates of *E. salina* shared the putative subset of eukaryotic homologs required for cholesterol biosynthesis, indicating cholesterol production is a shared feature of the species (Dataset S1).

**Fig 2.**
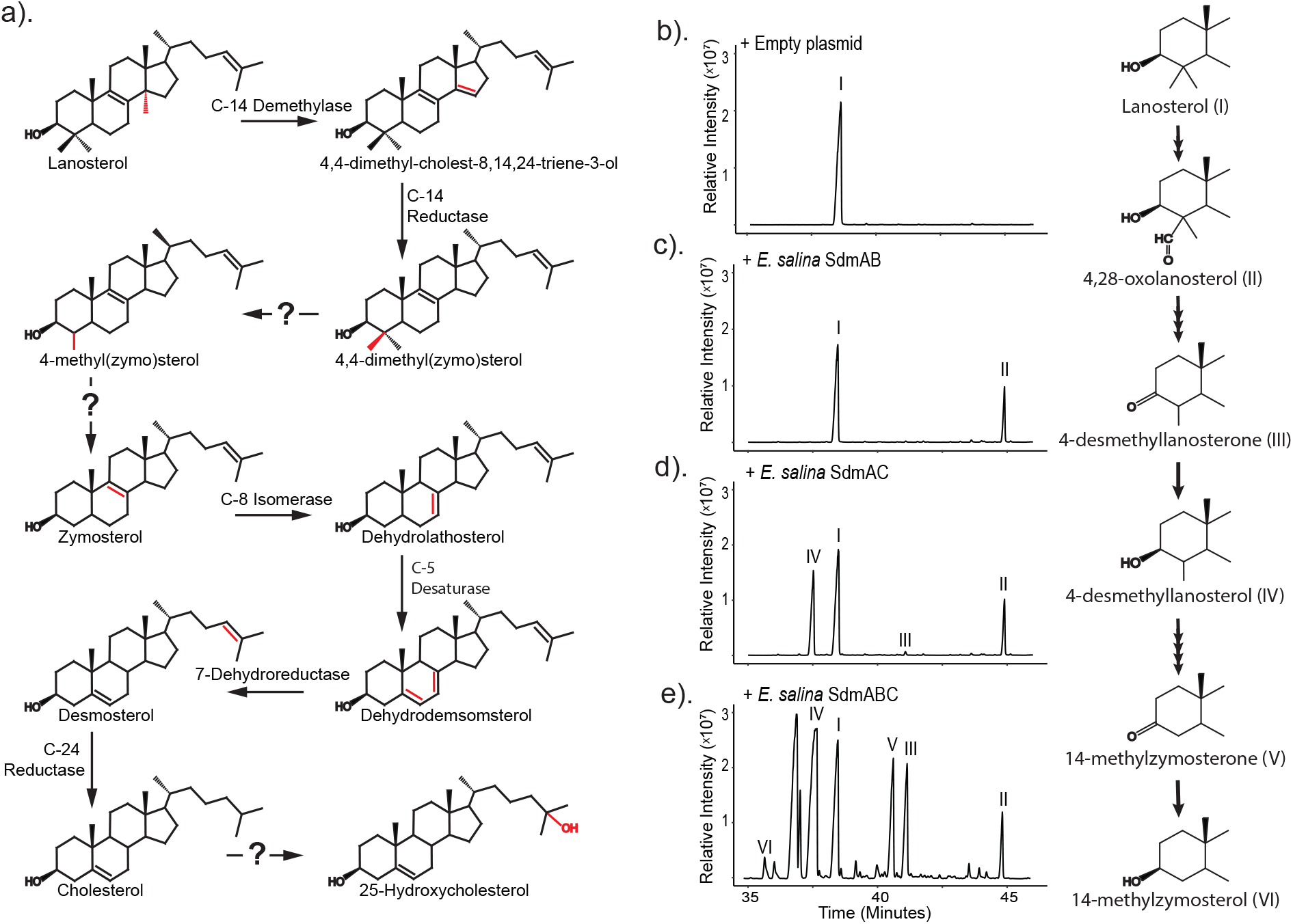
Sterol biosynthesis in *E. salina*. (a) Putative cholesterol biosynthesis pathway in *E. salina*. Homologs to the canonical eukaryotic proteins for cholesterol biosynthesis were identified by BLASTp search (<1 x e^-30^, 30% ID). *E. salina* does not have homologs to the eukaryotic enzymes responsible for C-4 demethylation and C-25 hydroxylation. Bacterial enzymes for complete C-4 sterol demethylation and C-25 hydroxylation have not been identified. Genomic context for identified *E. salina* genes are shown in SI Appendix, Fig. S5. (b) Total ion chromatograms showing lanosterol (I) substrate production in our *E. coli* heterologous expression system. (c) Co-expression of SdmAB homologs from *E. salina*, resulting in production of only a 4-methylaldehyde intermediate (III). These two enzymes are insufficient to demethylate at C-4, distinguishing them from SdmAB homologs in aerobic methanotrophs. (d) Co-expression of SdmAC homologs from *E. salina*, resulting in removal of single methyl group at C-4. (e) Co-expression of SdmABC homologs from *E. salina*, resulting complete demethylation at C-4. Lipids were derivatized to trimethylsilyl groups. C-4 demethylation intermediates were confirmed in a previous study and identified here by comparison to published mass spectra (32). Mass spectra of identified sterols are shown in *SI Appendix*, Fig. S7.

### C-4 demethylation in *E. salina* is distinct from eukaryotes and aerobic methanotrophs

While *E. salina* shares much of its cholesterol biosynthesis pathway with eukaryotes, it does not have homologs to the three eukaryotic proteins responsible for C-4 demethylation. Instead, *E. salina* has homologs to the dioxygenase-reductase pair, SdmAB, used by aerobic methanotrophs to remove only a single methyl group at C-4 (32). To test whether the *E. salina* SdmAB homologs are sufficient to remove both methyl groups at C-4 as required to produce cholesterol, we employed a heterologous expression system in *E. coli* engineered to overproduce lanosterol (Fig 2b)(34). Expression of the SdmA homolog alone in presence of lanosterol resulted in production of a 4-methylaldehyde intermediate. Expression of the SdmB homolog alone did not generate any demethylation intermediates (*SI Appendix*, Fig. S7 and Fig. S8). Interestingly, co-expression of the SdmAB homologs did not result in removal of any methyl groups at C-4 (Fig 2c), distinguishing SdmAB in *E. salina* from the homologs found in aerobic methanotrophs and indicating these two proteins are not sufficient to fully demethylate at the C-4 position in *E. salina*.

We next sought to determine whether the lack of C-4 demethylation by the *E. salina* homologs was tied to SdmA or SdmB. To do so, we leveraged the SdmA and SdmB homologs from *Methylococcus capsulatus* - which together remove a single methyl group in our expression system - and expressed them with their reciprocal SdmA or SdmB partner from *E. salina*. Expression of SdmA from *E. salina* with SdmB from *M. capsulatus* resulted in removal of a single methyl group at C-4, while expression of SdmB from *E. salina* with SdmA from *M. capsulatus* resulted in no demethylation at C-4 (*SI Appendix*, Fig. S9). Thus, we propose the *E. salina* SdmA homolog can carry out the oxygenation reactions required to demethylate, but the *E. salina* SdmB homolog is insufficient to carry out both the decarboxylation and reduction reactions required for demethylation at the C-4 position (32). These results led us to reassess *E. salina* for enzymes with the potential to carry out the decarboxylation and reduction reactions required for C-4 demethylation.

Through an additional BLASTp search of the *E. salina* genome, we identified another SDR-type reductase homologous to SdmB, SdmC (1 x e^-120^, 51% identity), restricted to the Myxococcota suborder Nannocystaceae (*SI Appendix*, Fig. S10). Expression of SdmC alone did not result in accumulation of any demethylation intermediates (*SI Appendix*, Fig. S8). However, co-expression of SdmC with SdmA from *E. salina* results in removal of a single methyl group at C-4, suggesting SdmC is capable of decarboxylating the oxidized methyl group and reducing the remaining ketone at C-3 into a hydroxyl as was shown for the *M. capsulatus* SdmB (Fig. 2c)(32). Additionally, co-expression of SdmC with *E. salina* SdmAB results in removal of both methyl groups at C-4 and production of additional demethylation intermediates (Fig. 2d). Thus, full demethylation at the C-4 position in *E. salina* is distinct from both bacterial C-4 demethylation in aerobic methanotrophs and from eukaryotic C-4 demethylation (*SI Appendix*, Fig. S11).

### *Calothrix* sp. NIES-4105 synthesizes conjugated sterols

Phylogenomic studies have identified several bacteria with the genomic capacity to produce complex sterols in phyla where sterol biosynthesis is underexplored (27)(28). The cyanobacterium *Calothrix* sp. NIES-4105 harbors a putative biosynthetic gene cluster that includes homologs for complex sterol biosynthesis, prompting us to analyze sterols in *Calothrix*. We were unable to detect any free sterols from Bligh-Dyer extractions (Fig 3a), distinguishing sterol production in *Calothrix* from *E. salina*. However, by acid hydrolyzing cell biomass, we detected sterols in the released ester and ether bound lipids (Fig 3b, *SI Appendix*, Fig. S1). Released sterols included cholesterol, representing a second instance of bacterial cholesterol biosynthesis. We identified several C-24 unsaturated intermediates associated with the Bloch biosynthesis pathway, including zymosterol, dehydrolathosterol and desmosterol. We also identified lathosterol, a C-24 saturated intermediate typical of the Kandutsch-Russell (K-R) cholesterol pathway. This combination of intermediates suggests *Calothrix* performs C-24 reduction as both an intermediary and final step in cholesterol synthesis, using both the Bloch and K-R biosynthesis pathways. Additionally, we identified 25-OHC in hydrolyzed *Calothrix* extracts, providing further evidence that *Calothrix*, like *E. salina*, produces conjugated sterols. Finally, to evaluate potential sterol contamination, we solvent extracted *Calothrix* culturing medium without bacterial biomass which we found to be devoid of sterols (*SI Appendix*, Fig. S3).

**Fig 3.**
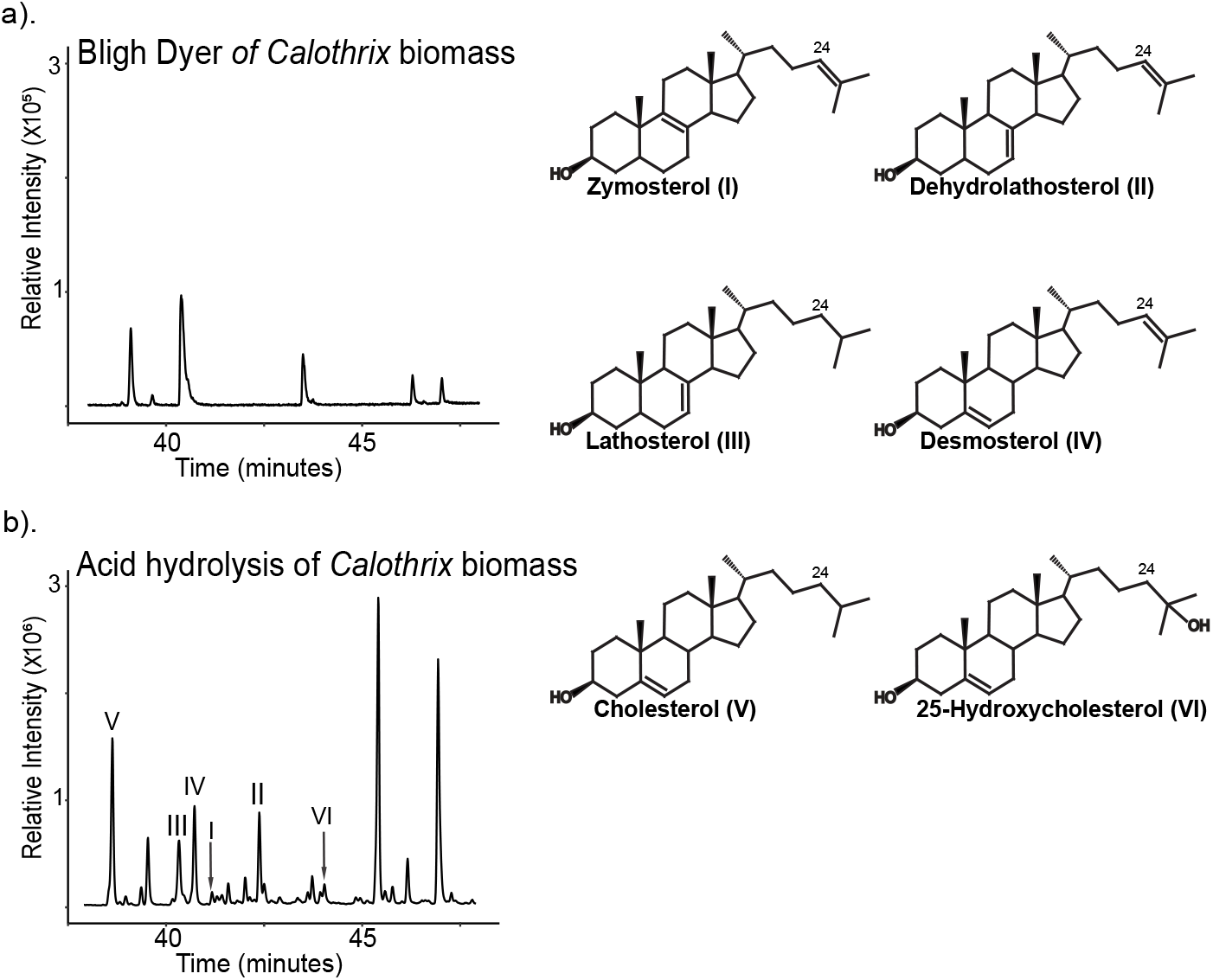
*Calothrix* synthesizes bound sterols. (a) Total ion chromatograms of free lipids extracted from *Calothrix* using a modified Bligh Dyer procedure. Peaks present in this chromatogram were not sterols based on retention time and comparison to a sterol standard mix. (b) Total ion chromatogram of ether and ester bound lipids extracted from *Calothrix* biomass. Sterol identified included cholesterol (V) and both C-24 saturated (III) and unsaturated intermediates (I, II, and IV). Additionally, hydrolysis released 25-hydroxycholesterol (VI). All lipids were derivatized to trimethylsilyl groups. Mass spectra of identified sterols are shown in *SI Appendix*, Fig. S1.

Throughout the course of lipid analyses, *Calothrix* was serial passaged and ceased sterol production. Changes in bacterial phenotype over continuous laboratory culture is a well-established occurrence and can be caused by many factors, including genetic mutation or changes in transcriptional or translational regulation (35). To determine if loss of sterol production in *Calothrix* was due to a genetic mutation, we sequenced the genome of the serial passaged strain and compared it to the reference genome of *Calothrix* sp. NIES-4105. We detected several mutations in the serial passaged strain (*SI Appendix*, Table S2). However, none of these mutations occurred in or upstream of putative sterol biosynthesis genes or in potential transcriptional regulators, suggesting the loss of sterol production is unlikely to be caused by genetic mutation. Further, to confirm expression of sterol biosynthesis genes from the passaged strain results in sterol production, we expressed the *osc* homolog in a heterologous *E. coli* system resulting in lanosterol production (*SI Appendix*, Fig. S12)(36)(37). Taken together, these results demonstrate that though the serial passaged strain of *Calothrix* no longer produces sterols under laboratory conditions, it maintains the genetic potential for sterol production.

### Sterol Biosynthesis in *Calothrix* Diverges from the Eukaryotic Pathway

We conducted a BLASTp search (<1 x e^-30^, 30% ID) for homologs to both the eukaryotic and *E. salina* cholesterol biosynthesis pathway in the *Calothrix* genome (*SI Appendix*, Table S1). *Calothrix* shares homologs for initial steps in cholesterol biosynthesis including oxidosqualene cyclization, demethylation at C-14 and C-4, and isomerization at C-8 (Fig. 4a). These genes occur in a single 23kb cluster alongside other putative sterol biosynthesis genes (Fig. 4b). *Calothrix* does not have homologs to the proteins required for C-5 desaturation, C-7 reduction, and C-24 reduction. The closest predicted homologs in *Calothrix* to either the eukaryotic or *E. salina* sterol synthesis proteins are located outside the sterol biosynthesis cluster and are more homologous to annotated non-sterol biosynthesis genes in *E. salina*. Additionally, we conducted a BLASTp search (<1 x e^-50^, 30% ID) for other bacterial species with the subset of cholesterol biosynthesis genes identified in *Calothrix*. We found 19 bacteria harbored this subset of genes (Dataset S1), although we note few have been tested for the presence of sterol molecules.

**Fig 4.**
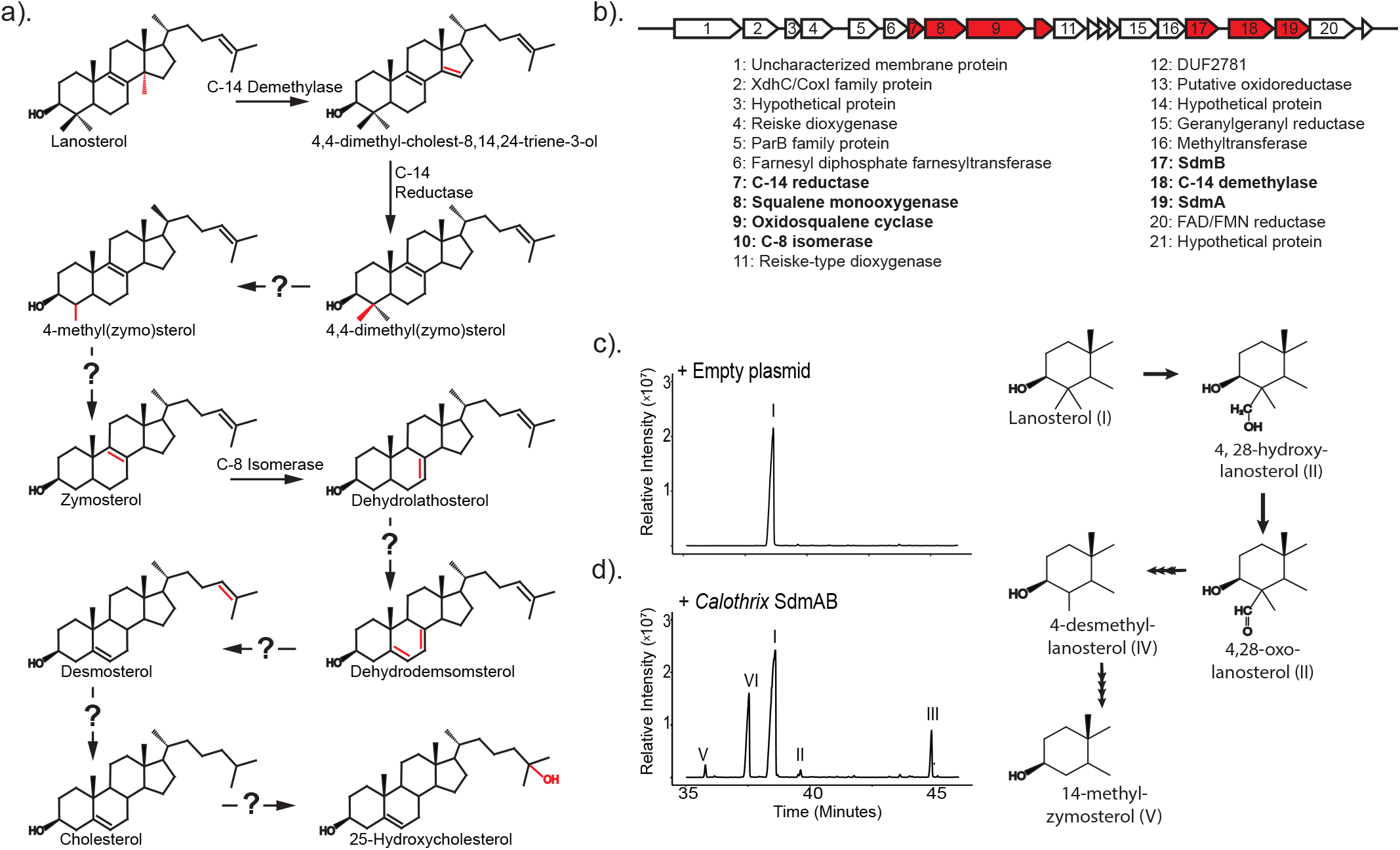
Sterol biosynthesis pathways in *Calothrix*. (a) Putative cholesterol biosynthesis pathway in *Calothrix*. Homologs to the canonical eukaryotic proteins for cholesterol biosynthesis were identified by BLASTp search (<1 x e^-30^, 30% ID). *Calothrix* does not have homologs to the eukaryotic enzymes responsible for C-4 demethylation, C-5 desaturation, C-7 reduction, C-24 reduction and C-25 hydroxylation. (b) Sterol biosynthesis genes in Calothrix are localized in a single gene cluster. Genes identified by our BLAST search are indicated in red and text labels bolded. Of note are several other genes annotated as putative biosynthesis genes which may be responsible for carrying out the missing steps in cholesterol biosynthesis. (c) Total ion chromatogram of lanosterol substrate production in our heterologous expression system. (d) Co-expression of SdmAB homologs from *Calothrix* showing complete demethylation at C-4. Lipids were derivatized to trimethylsilyl groups. C-4 demethylation intermediates were confirmed in a previous study and identified here by comparison to published mass spectra (32). Mass spectra of identified sterols are shown in *SI Appendix*, Fig. S2

*Calothrix* also utilizes SdmAB homologs to demethylate at C-4. Unlike *E. salina*, which requires two SDR-type reductases - SdmB and SdmC - for the removal of both methyl groups at C-4, *Calothrix* only harbors a single SdmB homolog, prompting us to test the biochemical capacity of the *Calothrix* SdmAB homologs (Fig 4c). Expression of the *Calothrix* SdmA homolog alone in the presence of lanosterol resulted in production of a 4-methylaldehyde intermediate. Expression of the SdmB homolog alone resulted in the production of lanosterone (*SI Appendix*, Fig S7 and S8). Co-expression of *Calothrix* SdmAB was sufficient to remove both methyl groups at C-4, demonstrating a third bacterial C-4 demethylation pathway, distinct from *E. salina*, aerobic methanotrophs and eukaryotes (*SI Appendix*, Fig S11).

## Discussion

In this study, we identified two bacteria capable of *de novo* cholesterol biosynthesis. The discovery of cholesterol and its intermediates demonstrates bacteria are capable of similar biosynthetic complexity associated with eukaryotic sterol synthesis and suggests nuanced regulatory and physiological roles for sterols in bacteria. This in part is exhibited by differences in pathway ordering. In eukaryotes, use of the Bloch or K-R pathway plays a regulatory role in lipid production and downstream metabolite synthesis (8)(7)(38). The intermediates we observed in *E. salina* and *Calothrix* suggest *E. salina* preferentially uses the Bloch pathway while *Calothrix* uses both the Bloch and K-R pathways. Use of both pathways by *Calothrix* may be indicative of varying functions for sterol intermediates and potentially novel regulatory systems driving the synthesis of these different intermediates. Further, the production of free versus conjugated sterols in bacteria provides another example of the potential complexity of sterol physiology in bacteria. Conjugations to other macromolecules impact the biophysical properties of sterols (39) and, in eukaryotic systems, these modified lipids are involved in specific functions including lipid storage and cell defense (19)(18). We demonstrated that both *E. salina* and *Calothrix* conjugate their sterols, but each do so to different extents - *E. salina* produces both free and conjugated sterols while *Calothrix* only produces conjugated sterols. However, given the difficulty in culturing of *Calothrix*, we cannot rule out free sterol production under different growth conditions. Regardless, these different sterol pools suggest a varied set of physiological roles for sterols in bacteria. Furthermore, bacterial sterol conjugation is not limited to *E. salina* and *Calothrix*; other bacteria conjugate sterols, either as the product of *de novo* biosynthesis (23) or modification of exogenously acquired sterols (40)(41). Expanding the extraction techniques used to analyze sterol lipids, identifying sterol conjugates present, and exploring enzymes responsible for downstream modifications would allow a better assessment of conjugated sterol diversity in the bacterial domain while providing a foundation for further exploration of sterol function and modifications.

Cholesterol production by *E. salina* and *Calothrix* also points to a complicated evolutionary history for sterol biosynthesis in the bacterial domain. Complex sterol biosynthesis is an ancient process; oxidosqualene cyclase (OSC), responsible for sterol cyclization, is thought to have evolved around the Great Oxidation Event (33) and many modern-day eukaryotic sterol biosynthesis genes are thought to predate the last eukaryotic common ancestor (6). Sterol biosynthesis in bacteria has been posited to be a product of horizontal gene transfer from eukaryotes, supported by its rarity and uneven distribution across bacterial phyla (33). However, recent phylogenetic and structural analyses of OSC and CYP51, responsible for sterol C-14 demethylation, suggest a bacterial origin for these proteins (28)(27)(42). The putative *E. salina* cholesterol biosynthesis pathway we identified is largely homologous to the eukaryotic pathway, indicating a shared evolutionary history for much of cholesterol biosynthesis between this myxobacterium and eukaryotes, although our analysis does not provide insight into the directionality of acquisition. We did find various cholesterol biosynthesis proteins, specifically those involved in desaturation modifications which have not been considered evolutionarily, occur in diverse bacterial phyla (Dataset S1). Further biochemical and phylogenetic analyses of these downstream genes may better resolve the evolutionary relationships governing sterol production in these two domains.

The putative *Calothrix* cholesterol biosynthesis pathway further diverges from the eukaryotic pathway, suggesting the existence of a potential novel cholesterol biosynthesis pathway. We posit *Calothrix* has evolved an independent mechanism to carry out desaturation at C-5 and saturation at C-7 and C-24. The *Calothrix* sterol biosynthesis gene cluster harbors several genes with annotated functions that suggest they may be able to perform the chemical reactions required to perform these steps. If that is the case, these additional instances of independent evolution towards cholesterol production highlight the versatility and importance of this lipid and further demonstrate the distinct evolutionary history underpinning sterol biosynthesis in bacteria. These missing homologs also raise questions about the biosynthetic capabilities of other bacteria. Because genes for desaturation modifications in *Calothrix* have not been identified, bioinformatic analysis alone is insufficient to identify other bacteria with the capacity of complex sterol production. We found the genes required for downstream biosynthesis, such as C-14 and C-4 demethylation, are present in a diverse set of bacteria including other myxobacteria and cyanobacteria, as well as actinobacteria, acidobacteria, and nitrospirea (Supp data file 1)(28). In many of these bacteria, sterol production has yet to be analyzed. Coupling more robust lipid analysis of these bacteria with further study of biosynthetic mechanisms would allow us to better grasp the distribution, diversity, and complexity of sterols in bacteria.

Finally, our study highlights how C-4 demethylation continues to provide a distinct example of independent evolution in sterol biosynthesis. We identified proteins responsible for complete C-4 demethylation in *E. salina* and *Calothrix* which function through a mechanism separate from eukaryotes, aerobic methanotrophs, and each other. In eukaryotes, C-4 demethylation is an oxygen dependent mechanism carried out by three proteins - a C-4 sterol methyl oxidase (ERG25/SMO), C-4 decarboxylase (ERG26/3β-HSD/D), and a 3-ketosteroid reductase (ERG27/3-SR) (43)(44)(45). All three proteins are involved in an iterative process that sequentially removes both methyl groups at the C-4 position although plants have been shown to utilize two distinct SMO proteins to remove each methyl group in a nonsequential manner (46). We have previously shown that in aerobic methanotrophs C-4 demethylation results in the removal of one methyl group and is carried out by a Rieske-type oxygenase, SdmA, and an NADP-dependent reductase, SdmB (32). Although both *E. salina* and *Calothrix* harbored homologs of SdmA and SdmB, it was unclear if these two proteins would be sufficient to remove both methyl group at the C-4 position. We show that the two *Calothrix* homologs are indeed sufficient to remove both methyl groups at C-4 but *E. salina* requires a second reductase to fully demethylate. These independent mechanisms for C-4 demethylation establish that cholesterol biosynthesis in these bacteria is not a simple case of horizontal gene transfer. Rather, it represents a case of convergent evolution in sterol biology that implies a critical role of C-4 sterol demethylation in sterol function in all domains of life. Indeed, C-4 demethylation has been shown to be required for proper function in eukaryotes as C-4 demethylase mutations are often lethal (47)(48)(43). What remains unclear, is what functional role C-4 demethylation plays in bacterial cells and if it is required for proper sterol function as observed in eukaryotes. Further biochemical and structural characterization of the various bacterial C-4 demethylation pathways should provide comparative insights into the functional significance of this modification.

## Materials and Methods

### Bacterial Culture

Strains used in this study are listed in *SI Appendix, Table S3. E. salina* DSM 15201 was cultured in artificial seawater liquid medium, supplemented with autoclaved and concentrated whole cell *E. coli*, at 30°C with shaking at 225 rpm (Thermo Scientific, MaxQ8000), for 14 days (29). *Calothrix* sp. NEIS-4105 was cultured in BG-11 liquid medium at 25°C for 60 days with 10-hour light, 14-hour dark cycles. Supplemental prey *E. coli* was cultured in LB at 37°C, shaking at 225rpm for 18hr, before concentrating 10:1 (vol:vol) in water and autoclaving. *E. coli* heterologous expression strains were cultured in TYGPN medium at 30°C or 37°C, shaking at 225 rpm and supplemented, if necessary, with gentamycin (15 μg/mL), kanamycin (30 μg/mL), carbenicillin (100 μg/mL), and/or chloramphenicol (20 μg/mL).

### Lipid Extraction and GC-MS Analysis

Lipid extraction protocol are described in detail in *SI Appendix, Methods*. Unbound sterols were extracted from cell pellets through a modified Bligh-Dyer extraction (30). Where applicable, sterols extracted through the Bligh-Dyer extraction were further separated into fractions based on polarity using Si-gel chromatography (49). Bound sterols were analyzed by hydrolyzing either lipid extracts, Si-gel chromatography fractions, or lyophilized cell pellets in methanolic base or acid. All lipids were derivatized to trimethylsilyl ethers with 1:1 (vol:vol) Bis(trimethylsilyl)trifluoroacetamide: pyridine before analysis on an Agilent 7890B Series GC using methods described in *SI Appendix, Methods*.

### Bioinformatic Analysis

To identify sterol biosynthesis genes in *E. salina* and *Calothrix*, we conducted a BLASTp search (50). To be considered a putative sterol biosynthesis gene, we set an e-value cut-off of 1 x e^-30^ and percent identity cutoff of 30. BLASTp search results are listed in *SI Appendix*, Table S1. To identify other bacteria which the sterol biosynthesis genes we identified in *E. salina* and *Calothrix*, we first conducted a BLASTp search of bacteria in the genomic databases on the JGI IMG portal for OSC (<1 x e ^-50,^ 30%ID). Of this subset of bacteria, we conducted further BLASTp searches (<1 x e ^-50,^ 30%ID) using identified proteins from *E. salina* and *Calothrix* (51)(52). Phylogenetic analyses of SdmBC homologs are further described in *SI Appendix, Methods*.

### Genomic Sequencing and Mutation Identification

Genomic DNA from *Calothrix* sp. NIES-4105 was isolated through phenol-chloroform extraction. Library preparation (Illumina DNA Prep Kit; San Diego, CA) and whole genome sequencing was performed by SeqCenter (SeqCenter; Pittsburgh, PA) and sequenced on an Illumina NextSeq 2000, producing 2×151bp reads. Bcl-convert (v3.9.3) was used for demultiplexing, quality control, and trimming. To identify mutations in the serial passaged strain, variant calling was performed using breseq (v0.35.4)(53).

### Molecular Cloning

Cloning protocols are described in *SI Appendix, Methods*. Plasmids and oligonucleotides used in this study are described in *SI Appendix, Tables S4* and *S5*.

### Heterologous Expression

Sterol biosynthesis genes were overexpressed in *E. coli* from compatible plasmids with either an IPTG-inducible *lac* or arabinose-inducible *araBAD* promoters (34). Heterologous expression strains were constructed as describes in *SI Appendix, Table S6*. Genes of interest were expressed from the IPTG-inducible plasmid pSRKGm-*lac*UV5-rbs5 and/or the arabinose-inducible plasmid pBAD1031K. *E. coli* strains were cultured at 37°C, in 20 mL TYGPN medium supplemented with antibiotics (as necessary) until mid-exponential phase when expression was induced with 500 μM IPTG and 0.2% arabinose for 30-40 h at 30°C with shaking at 225 rpm, before harvest of cells.

## Supporting information

Data Set

## Acknowledgements

We thank members of the Welander lab for helpful discussions. Funding for this study was provided by National Science Foundation Award 1919153 (to P.V.W).

