## Supplemental File for "De novo cholesterol biosynthesis in the bacterial domain"

### Supporting Information for
De novo cholesterol biosynthesis in the bacterial domain

Alysha K. Lee, Jeremy H. Wei, and Paula V. Welander

Department of Earth System Science, Stanford University, Stanford, CA 94305

#### Supplementary Information Text

#### Methods

**Lipid Extraction.** Unbound sterols were extracted from lyophilized cell pellets using a modified Bligh-Dyer extraction protocol (1). Pellets were sonicated for 1 hour in 10:5:4 (vol: vol: vol) methanol: dichloromethane (DCM): water. Lipids were then phase separated using two times the volume 1:1 (vol: vol) DCM: water. The organic phase was transferred and evaporated under $N_2$ yielding total lipid extracts (TLE). Where indicated, TLE was fractionated by polarity using Si chromatography (2) with a column scheme as follows: 1.5 column volumes of hexane, 2 column volumes of 8:2 (vol: vol) hexane: DCM, 2 column volumes of DCM, 2 column volumes 1:1 (vol:vol) DCM: ethyl acetate, 2 columns volumes ethyl acetate yielding a alkynes, non-polar, ketone, alcohol, and polar fraction, respectively. To analyze bound sterol, lyophilized cell pellets, TLEs, or Si chromatography fractions were resuspended in 1N HCl or KOH in methanol and heated at 75C for 3 hours. Reactions were neutralized using KOH or HCl respectively and phases separated using twice the volume of 1:1 (vol: vol) DCM: water. The organic phase was transferred and evaporated under $N_2$.

**GC-MS Methods**. Lipids were separated on a 60m Agilent DB17HT column (60 m x 0.25 mm i.d. x 0.1 um film thickness) with helium as the carrier gas at constant flow of 1.1 mL/min and programed as follows: 100 $^{o}$C for 2 min; then 8 $^{o}$C/min to 250 $^{o}$C and held for 10 min; then 3 $^{o}$C/min to 330 $^{o}$C and held for 17 min. 2 μL of each sample was injected in splitless mode at 250 $^{o}$C. The GC was coupled to a 5977A Series MSD with the ion source at 230 $^{o}$C and operated at 70 eV in

EI mode scanning from 50-850 Da in 0.5 s. Lipids were identified based on retention time and comparison to previously confirmed laboratory standards, published spectra (3) (4) and spectra deposited in the American Oil Chemists' Society (AOCS) Lipid Library (http://lipidlibrary.aocs.org/index.cfm) and the National Institute of Standards and Technology (NIST) databases.

**Molecular Biology Techniques.** Oligonucleotides were purchased from Integrated DNA Technologies (Coralville, IA). Genomic DNA from *E. salina* was isolated using the GeneJET Genomic DNA Purification Kit (Thermo Scientific). Genomic DNA from *Calothrix* sp. NEIS-4105 was isolated using a phenol-chloroform extraction as previously described (5). Plasmid DNA was isolated using the GeneJET Plasmid Miniprep Kit (Thermo Scientific). DNA fragments used during cloning were isolated using the GeneJET Gel Extraction Kit (Thermo Scientific). DNA was sequenced by ELIM Biopharm (Hayward, CA).

**Plasmid Cloning**. Plasmids were constructed by sequence and ligation independent cloning (SLIC), adapted from (6). Briefly, complementary overhangs were created on gel-purified PCR product inserts and a restriction enzyme-linearized vector by incubation with T4 DNA polymerase (EMD Milipore) in the absence of nucleotides followed by annealing and transformation without ligation. *E. coli* strains were transformed by electroporation using a MicroPulser Electroporator (BioRad) as recommended by the manufacturer.

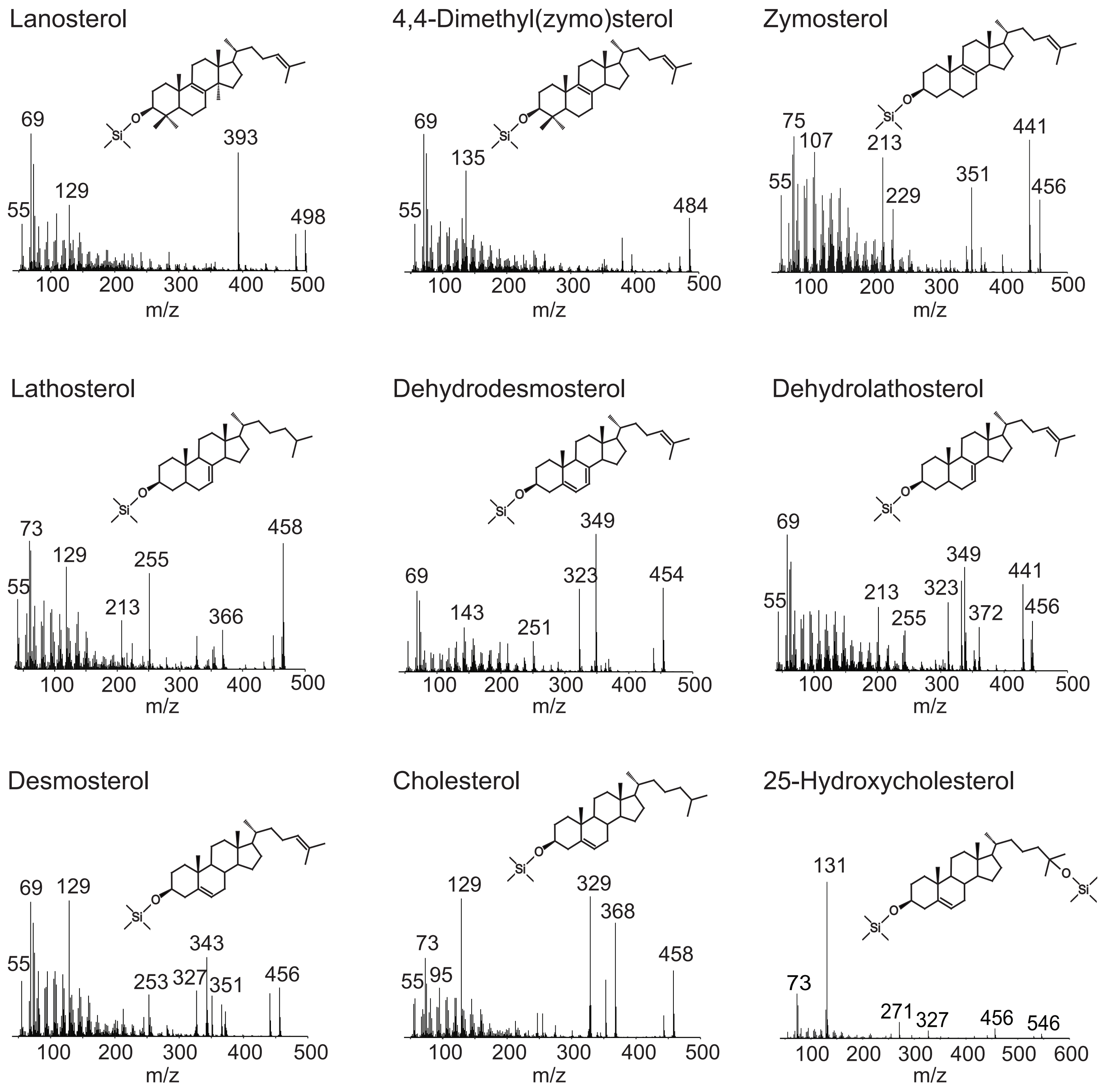

**Fig S1.** Mass spectra of sterols identified in *E. salina* or *Calothrix* NIES-4105 extracts. Extracted sterols were derivatized to trimethylsilyl (TMS) groups and separated on an Agilent 7890B series GC through a 60m Agilent DB17 column (60m x 0.25 mm i.d x 0.1 μm film thickness) with helium as the carrier gas coupled to a 5977A series MS. See Supplementary Methods section for full GC-MS method details.

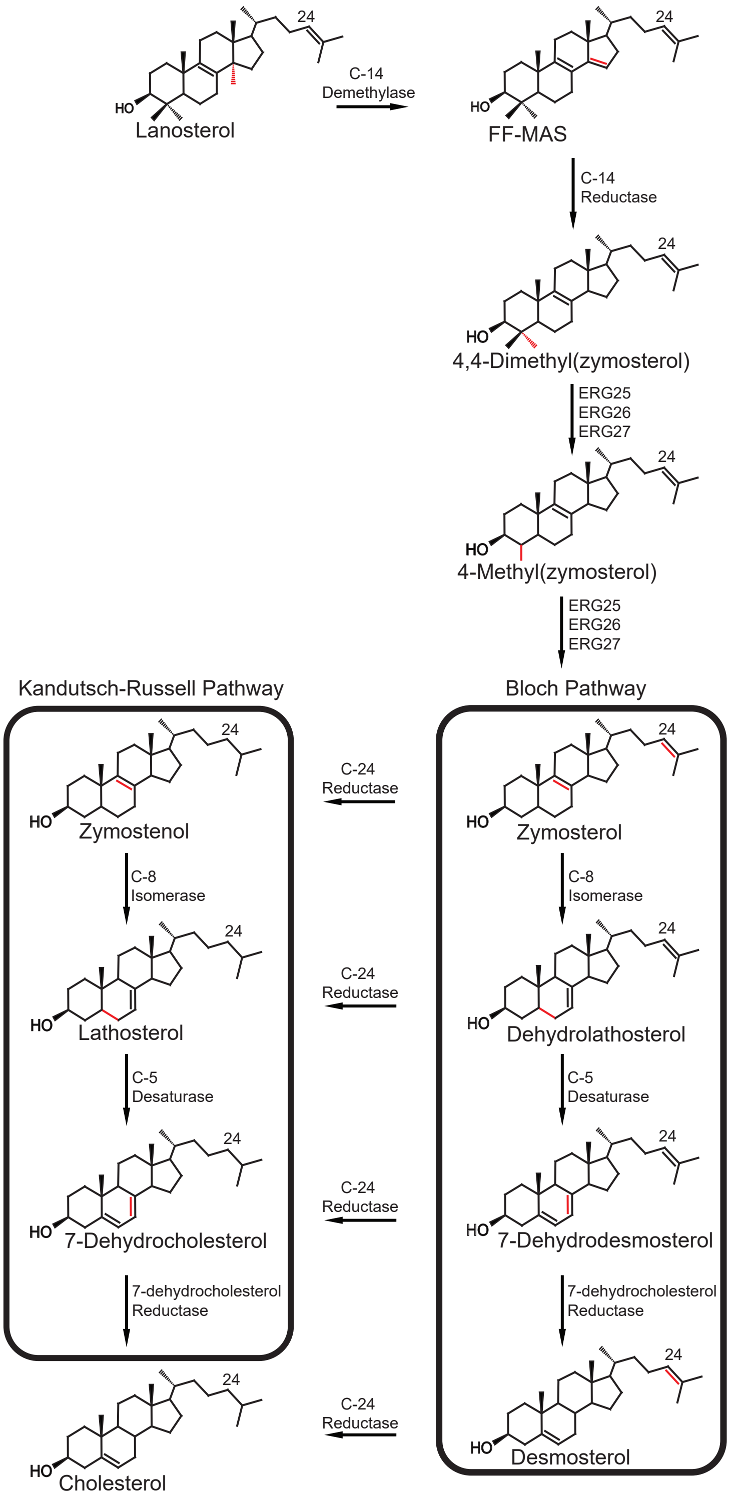

**Fig S2.** The Bloch versus Kandutsch-Russell (K-R) cholesterol biosynthesis pathways. The Bloch pathway proceeds from C-14 and C-4 demethylation through several steps modifying desaturation in the ring structure before saturation at C-24 as the final step. The K-R pathway utilizes the same enzymes to perform cholesterol biosynthesis but C-24 saturation occurs as an intermediary step. Recent work has suggested this likely occurs after C-4 and C-14 demethylation in eukaryotes (7).

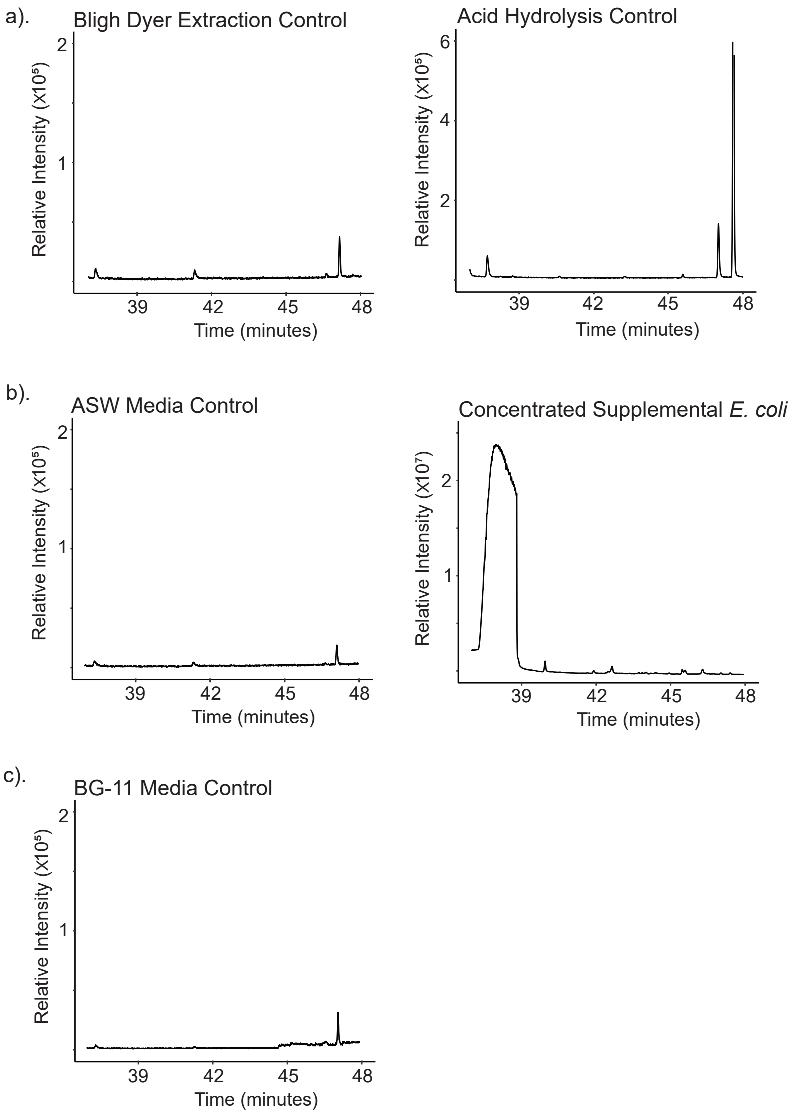

**Fig S3.** Total ion chromatograms of extraction controls and growth media. Total ion chromatogram of total lipid extracts (TLE) from (a) extraction processes used to analyze sterols in *E. salina* and *Calothrix*. Sterile water was added in place of bacterial biomass. (b) Extractions of media components used to culture *E. salina*. Both medium and supplemental *E. coli* were extracted using a modified Bligh Dyer technique. Extracted *E. coli* was concentrated 5-fold from what was used to culture *E. salina*. (c) Extraction of medium used to culture Calothrix. Medium was extracted using a modified Bligh Dyer technique. Spectra in peaks of all chromatographs were compared to the NIST database and published spectra and none were found to be sterols.

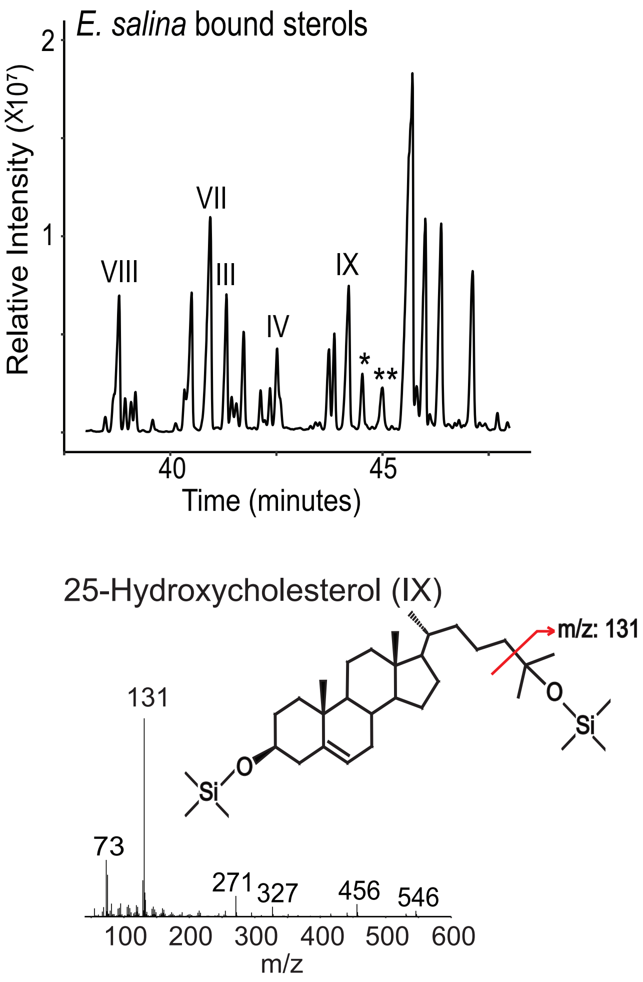

**Fig S4.** Probable hydroxysterols released from hydrolyzed *E. salina* lipid extracts. TMS-derivatized 25-hydroxycholesterol (25-OCH) has a diagnostic 131 peak that is produced from fragmentation of the tail. The presence of 25-OCH in *E. salina* samples was confirmed by comparing elution time to a known standard. Two neighboring peaks were found to share the same parent ion and diagnostic 131 peak, suggesting they may also represent sterols hydroxylated at C-25.

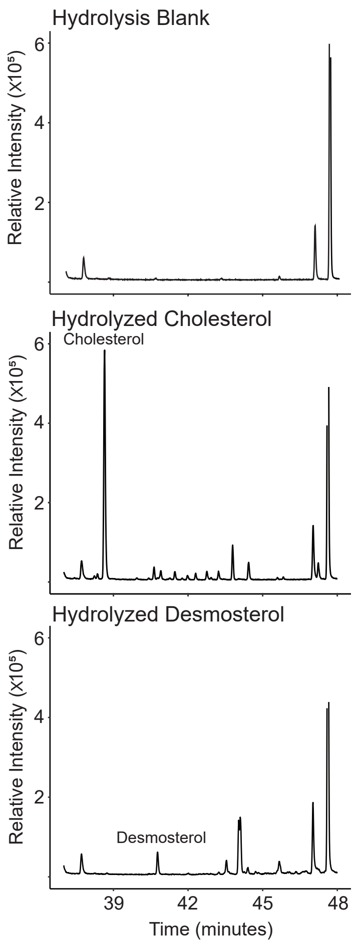

**Fig S5.** Acid hydrolysis of cholesterol and desmosterol standards. Total ion chromatograms of a hydrolysis blank, a hydrolyzed cholesterol standard, and a hydrolyzed desmosterol standard. Acid hydrolysis was performed as described in the Methods section. Lipids were derivatized to TMS groups. The spectra of peaks observed in the hydrolyzed standard chromatgrams were compared to a 25-hydroxycholesterol standard and spectra deposited in the NIST database, none were found to be oxysterols.

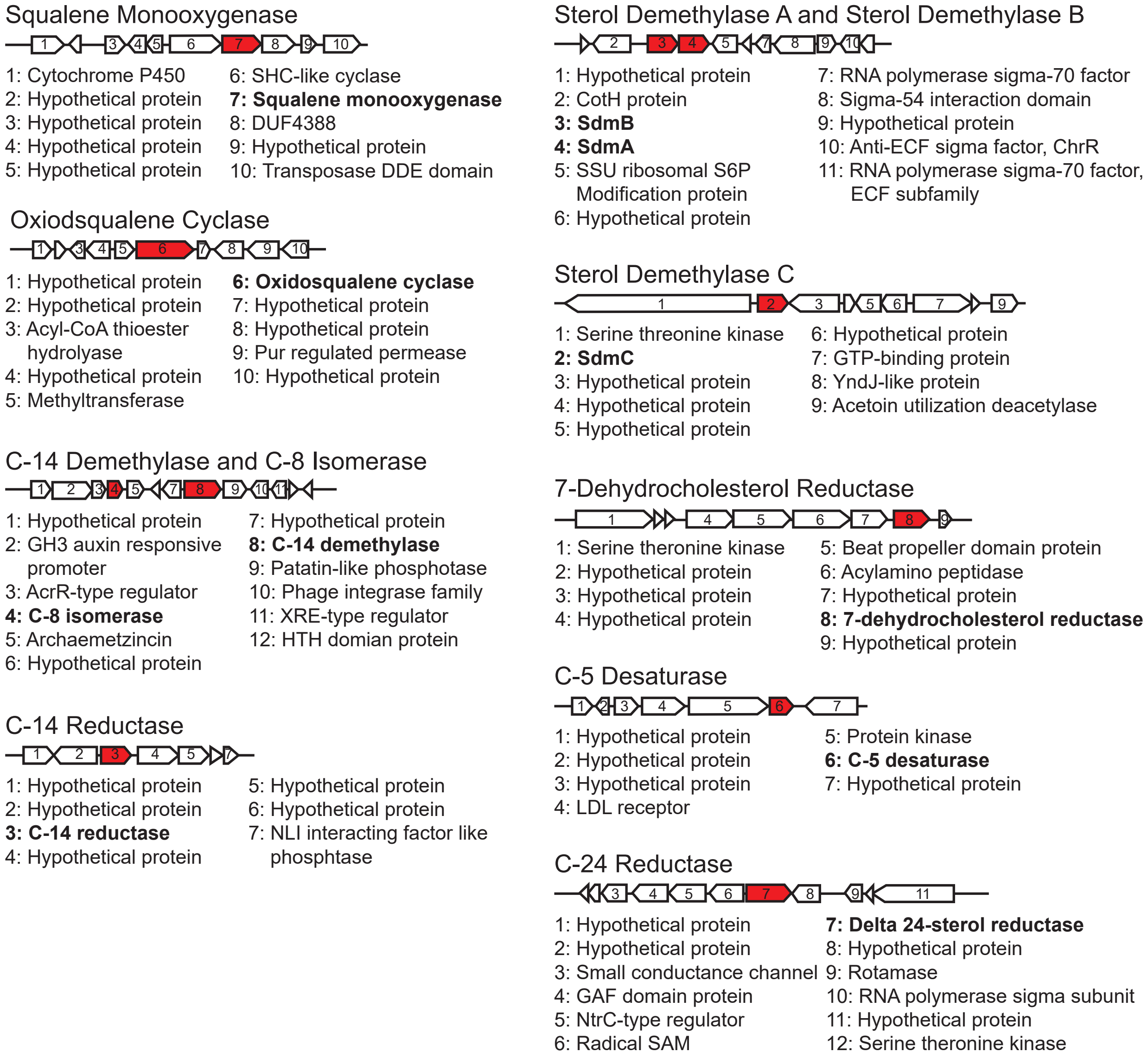

**Fig S6.** Genomic neighborhoods of sterol biosynthesis genes in *E. salina*. Sterol biosynthesis genes identified by our BLASTp search (<1xe $^{-30}$; 30% ID) are colored red and text labels bolded.

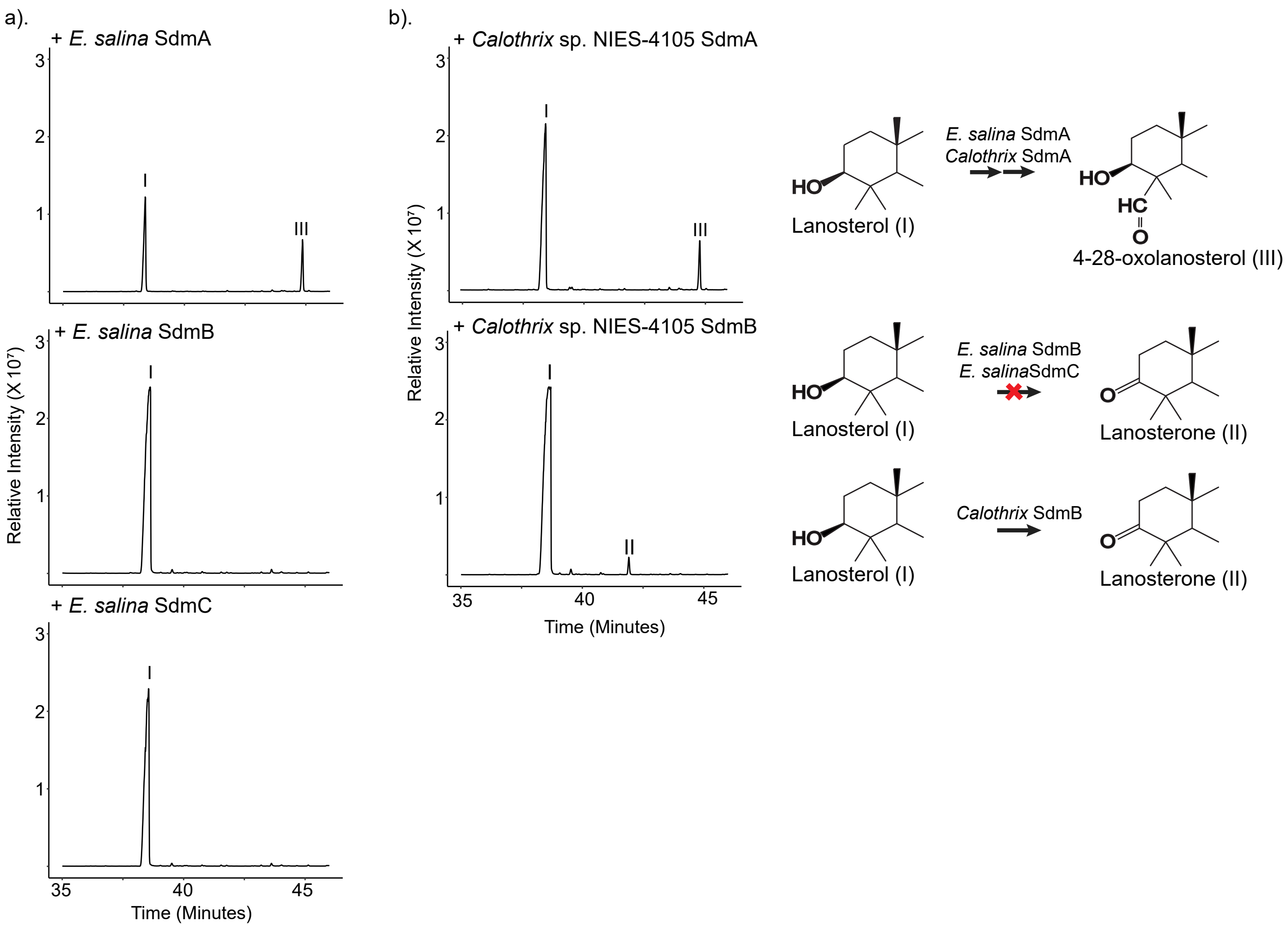

**Fig S7.** Total ion chromatograms of TLEs of (a) heterologous expression strains from *E. salina* SdmA, SdmB, and SdmC homologs and (b) heterologous expression strains from *Calothrix* SdmA and SdmB homologs. In both organisms, SdmA homologs produce the C-4 oxidation intermediate, 4-28-oxolanosterol. The SdmB and SdmC homologs from *E. salina* have no apparent effect on the substrate lanosterol. The SdmB homolog from *Calothrix* oxidizes the 3β-hydroxyl into a ketone, producing lanosterone.

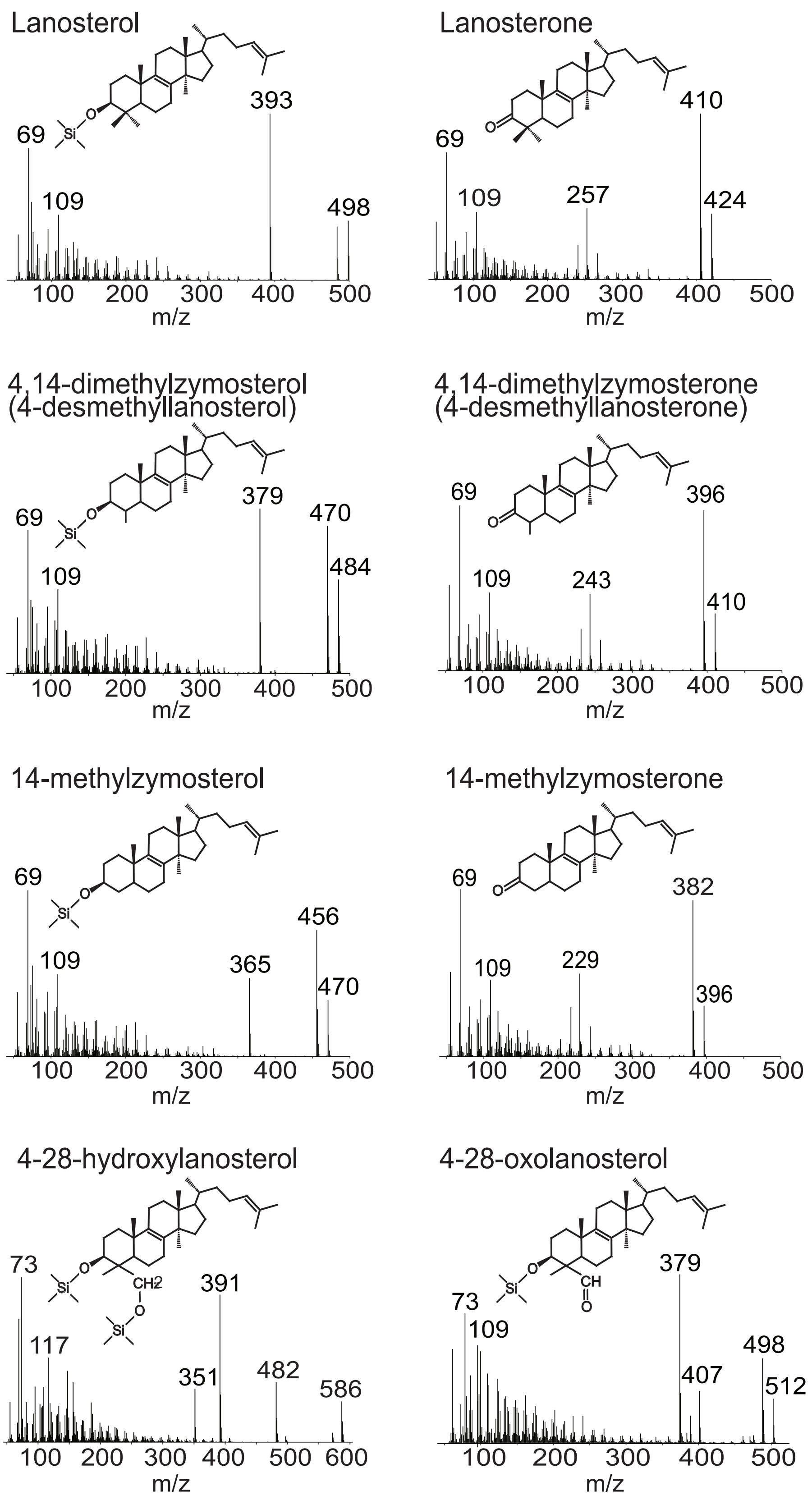

**Fig S8.** Mass spectra of sterols identified in heterologous expression cultures. Extracted sterols were derivatized to (TMS) groups and separated on an Agilent 7890B series GC with helium as the carrier gas and was coupled to a 5977A series MS. See Supplementary Methods section for full GC-MS method details.

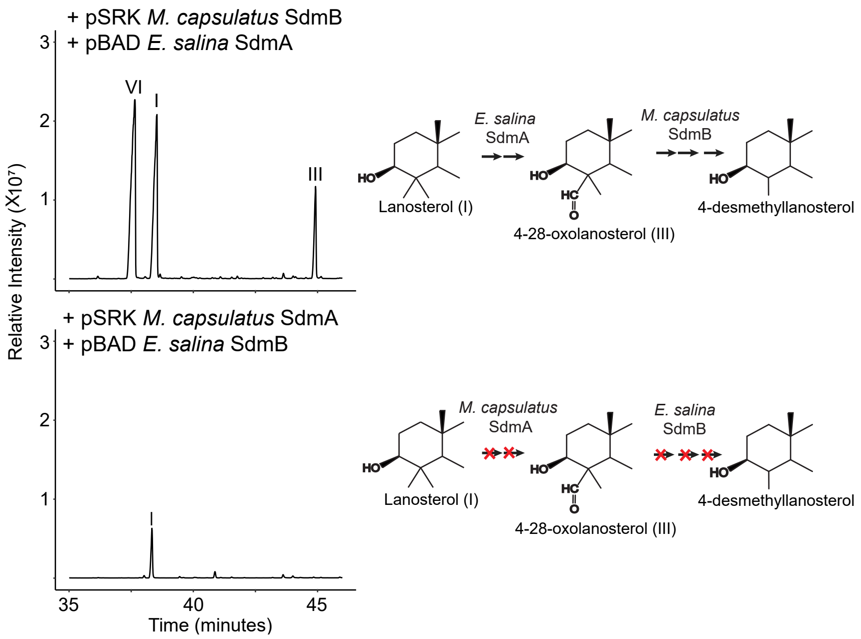

**Fig S9.** Heterologous expression of reciprocal SdmA and SdmB pairs from *E. salina* and *M. capsulatus*. SdmA and SdmB homologs form either *E. salina* or *M. capsulatus* were expressed from compatible plasmids in an *E. coli* strain engineered to overproduce lanosterol. Expression of *E. salina* SdmA with *M. capsulatus* SdmB resulted in single demethylation at C-4 while expression of *E. salina* SdmB with *M. capsulatus* SdmA did not, suggesting the *E. salina* SdmB homolog is insufficient to carry out the decarboxylation and reduction steps required to demethylate at C-4.

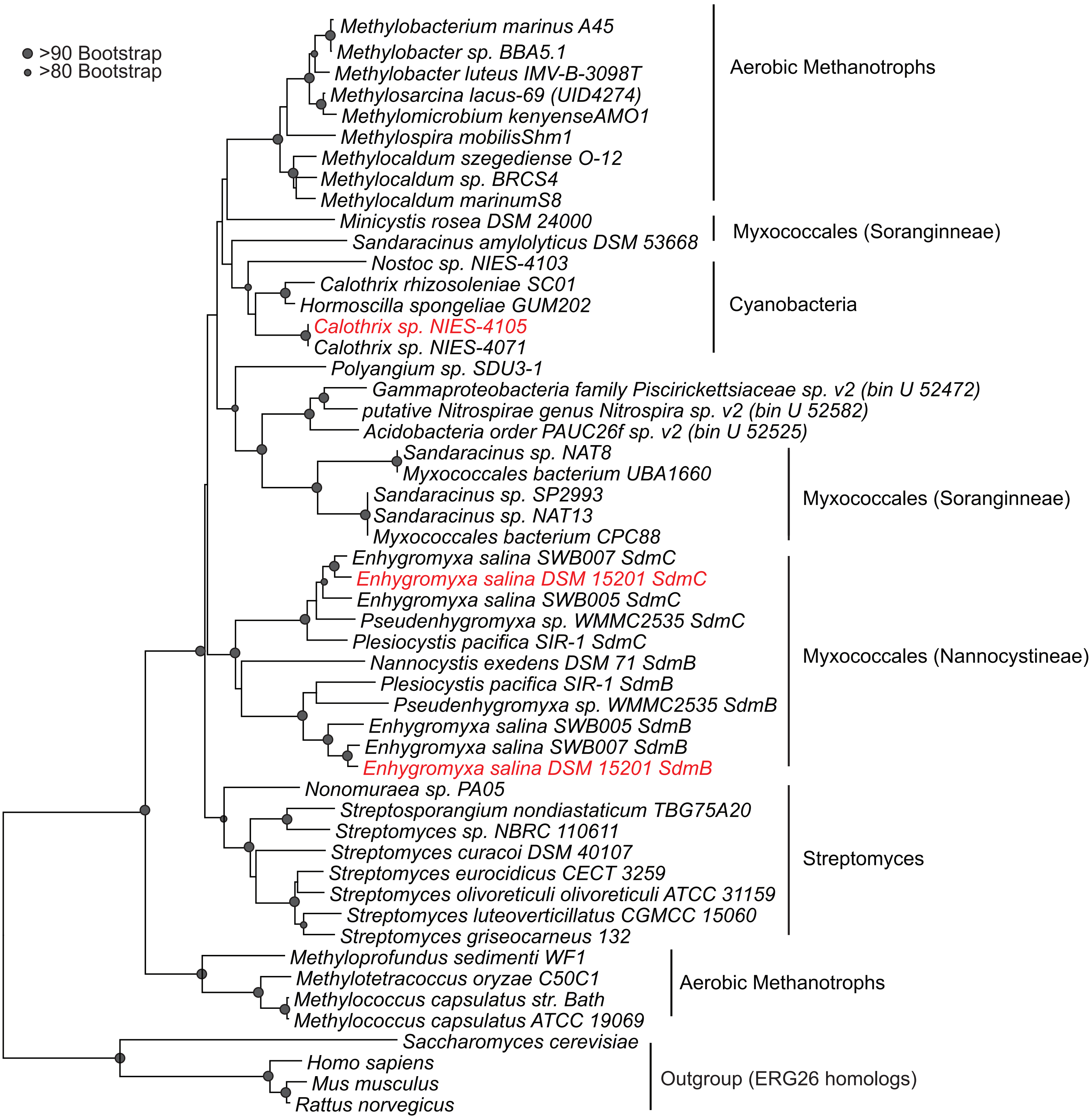

**Fig S10.** Neighbor-joining phylogenetic tree of SdmBC homologs. Homologs were identified by BLASTp search (e-value <1 x $e^{-50}$; 30% identity) of the genomic databases. Protein sequences were aligned by MUSCLE using MEGA. A neighbor-joining tree was generated using the gamma model, four gamma rate categories and 500 bootstrap replicates. SdmB homologs from *E. salina* and *Calothrix* and the SdmC homolog from *E. salina* tested in this paper are highlighted read. SdmC homologs are only present in the Myxococcata suborder Nannocystineae and form a monophyletic clade with SdmB homologs from the same suborder.

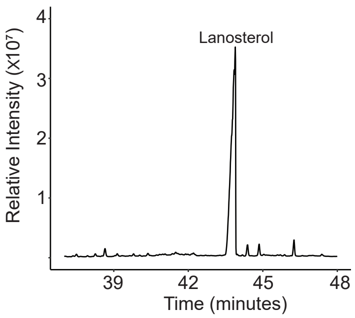

**Fig S12.** Total ion chromatogram of TLE from heterologous expression of oxidosqualene cyclase (*osc*) from serial passaged *Calothrix* strain. *osc* was amplified from genomic DNA extracted from serial passaged *Calothrix* strain, cloned into a compatible plasmid and expressed in an *E. coli* strain engineered to overproduce oxidosqualene. This resulted in production of lanosterol, demonstrating that the *osc* from the serial passaged *Calothrix* strain is functional.

a). C-4 demethylation in eukaryotes (vertebrates and fungi):

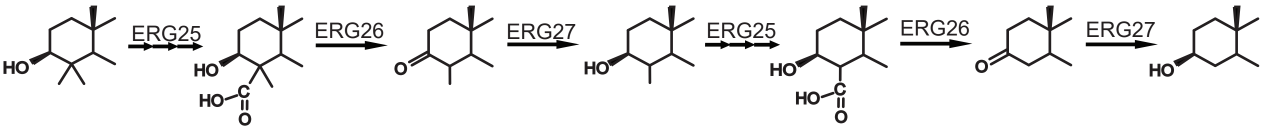

b). C-4 demethylation in aerobic methanotrophic bacteria:

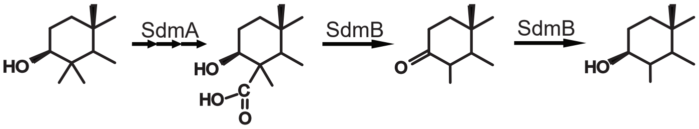

c). C-4 demethylation in *E. salina*:

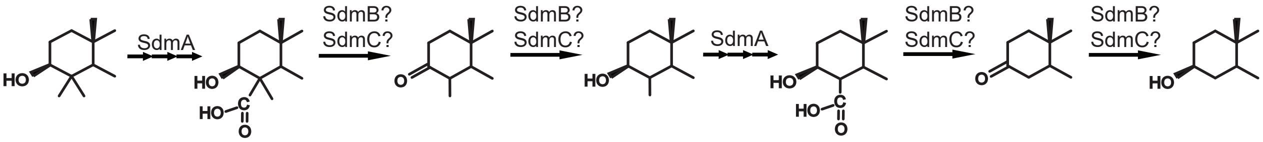

d). C-4 demethylation in *Calothrix*:

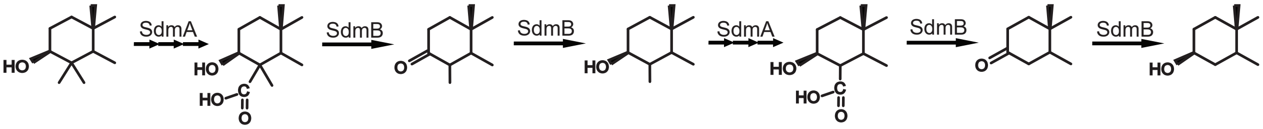

**Fig S11.** C-4 demethylation in eukaryotes and bacteria. a) In eukaryotes, C-4 demethylation is carried out by three enzymes. ERG25, a monooxygenase, performs three oxidation reactions to carboxylate the 4α-methyl group. ERG26, a short chain dehydrogenase reductase (SDR)-type reductase, catalyzes the decarboxylation of the 4α-methyl group and remaining methyl group epimerizes into the α-position. ERG27, an SDR-type reductase, reduces the C-3 ketone to a hydroxyl. This concerted reaction repeats a second time to remove the remaining methyl group. b) In aerobic methanotrophs, SdmA, a dioxygenase, performs the required oxidation reactions to carboxylate the 4β-methyl group. SdmB, an SDR-type reductase, then both decarboxylates the 4β-methyl group and reduces the C-3 ketone to a hydroxyl, producing the final monomethyl product. c) In *E. salina*, SdmA oxidizes one of the C-4 methyl groups to a carboxylate. SdmB and/or SdmC then perform the decarboxylation and reduction reactions. Further mechanistic study is required to untangle the role these two proteins have in C-4 demethylation. This reaction then repeats likely through the same mechanism. d) In *Calothrix*, SdmA oxidizes one of the methyl groups to a carboxylate. SdmB then performs the decarboxylation and reduction reactions to remove the first methyl group. This reaction then repeats, likely through the same mechanism.

**Table S1.** *E. salina* and *Calothrix* sp. NIES-4105 closest homologs to canonical sterol biosynthesis genes.

| | ***H. sapiens*** | ***S. cerevisiae*** | ***M. capsulatus*** | ***E. salina*** | ***Calothrix* sp. NIES-4105** |
|---|---|---|---|---|---|
| ***E. salina*** | | | | | |
| Squalene monooxygenase (Ga0097779_102654) | 1e$^{-27}$/25% | 2e$^{-28}$/24% | **2e$^{-37}$/29%** | N/A | **3e$^{-40}$/31%** |
| Oxidosqualene cyclase (Ga0097779_114516) | **6e$^{-112}$/33%** | **3e$^{-96}$/30%** | **1e$^{-169}$/44%** | N/A | **8e$^{-155}$/38%** |
| C-14 demethylase (Ga0097779_103211) | **4e$^{-97}$/36%** | **7e$^{-86}$/34%** | **8e$^{-133}$/44%** | N/A | **1e$^{-154}$/48%** |
| C-14 reductase (Ga0097779_105926) | **1e$^{-42}$/32%** | **2e$^{-41}$/29%** | N/A | N/A | **5e$^{-44}$/63%** |
| SdmA (Ga0097779_103152) | N/A | N/A | **3e$^{-104}$/45%** | N/A | **1e$^{-163}$/60%** |
| SdmB (Ga0097779_103151) | N/A | N/A | **2e$^{-79}$/41%** | N/A | **1e$^{-121}$/50%** |
| ERG25 (Ga0097779_103191) | 2e$^{-08}$/24% | 2e$^{-15}$/26% | N/A | N/A | 5e$^{-27}$/30% |
| ERG26 (Ga0097779_105318) | **2e$^{-46}$/32%** | **1e$^{-39}$/32%** | N/A | N/A | **3e$^{-44}$/31%** |
| ERG27 (Ga0097779_11785) | 8e$^{-12}$/26% | --- | N/A | N/A | 4e$^{-11}$/27%* |
| C-8 sterol isomerase (Ga0097779_103215) | **2e$^{-37}$/39%** | **8e$^{-47}$/43%** | N/A | N/A | **2e$^{-57}$/51%** |
| C-5 desaturase (Ga0097779_11078) | **1e$^{-45}$/35%** | **1e$^{-41}$/36%** | N/A | N/A | 4e$^{-16}$/30%* |
| 7-dehydrocholesterol reductase (Ga0097779_100662) | **1e$^{-87}$/36%** | --- | N/A | N/A | 2e$^{-16}$/30%* |
| Delta 24 sterol reductase (Ga0097779_101115) | **2e$^{-127}$/41%** | --- | **2e$^{-94}$/38%** | N/A | --- |
| ***Calothrix* sp. NIES-4105** | | | | | |
| Squalene monooxygenase (Ga0263810_115391) | 3e$^{-27}$/25% | 1e$^{-28}$/27% | **3e$^{-32}$/26%** | **2e$^{-40}$/31%** | N/A |
| Oxidosqualene cyclase (Ga0263810_115390) | **2e$^{-163}$/39%** | **3e$^{-105}$/32%** | **0.0/44%** | **9e$^{-155}$/38%** | N/A |
| C-14 demethylase (Ga0263810_115381) | **4e$^{-111}$/38%** | **1e$^{-78}$/33%** | **e$^{-131}$/44%** | **2e$^{-158}$/48%** | N/A |
| C-14 reductase (Ga0263810_115392) | 2e-$^{22}$/40% | 2e$^{-15}$/36% | N/A | **2e$^{-42}$/63%** | N/A |
| SdmA (Ga0263810_115380) | N/A | N/A | **1e$^{-102}$/44%** | **1e$^{-163}$/60%** | N/A |

| | | | | | |
|---|---|---|---|---|---|
| SdmB (Ga0263810_115382) | N/A | N/A | **7e$^{-97}$/45%** | **1e$^{-121}$/50%** | N/A |
| ERG25 (Ga0263810_118348) | 3e$^{-15}$/32% | 3e$^{-18}$/35% | N/A | 1e$^{-28}$/30%* | N/A |
| ERG26 (Ga0263810_118461) | **7e$^{-41}$/31%** | **8e$^{-33}$/30%** | N/A | 3e$^{-44}$/31%* | N/A |
| ERG27 (Ga0263810_119321) | 2e$^{-16}$/25% | | N/A | 2e$^{-13}$/27% | N/A |
| C-8 sterol isomerase (Ga0263810_115389) | **3e$^{-31}$/35%** | 5e$^{-27}$/36% | N/A | **2e$^{-57}$/49%** | N/A |
| C-5 desaturase (Ga0263810_118508) | 3e$^{-23}$/36% | 4e$^{-08}$/25% | N/A | 1e$^{-21}$/29% | N/A |
| 7-dehydrocholesterol reductase (Ga0263810_115392) | 2e$^{-16}$/31% | --- | N/A | 6e$^{-14}$/30% | N/A |
| Delta 24 sterol reductase (Ga0263810_116277) | 2e$^{-10}$/34% | ---- | 8e$^{-11}$/31% | 6e$^{-04}$/25% | N/A |

Sterol biosynthesis genes from *Homo sapiens*, *Saccharomyces cerevisiae*, and *Methylococcus capsulatus* were used as a query for BLASTp search (e-value < 1 x e$^{0.0}$) to identify putative cholesterol biosynthesis pathways in *E. salina* and *Calothrix*. Putative sterol biosynthesis genes in each organism were than used as a query for BLASTp search in the other bacterium. IMG database locus tags for top hits in each bacterium, with corresponding e-values and identities, are listed for each gene in the canonical cholesterol biosynthesis pathway. E-values colored green and bolded within our e-value cutoff (e-value < 1 x e$^{-30}$). E-values colored yellow and marked with an asterisk denote homologs where lowest e-value hit in either Calothrix or E. salina was not the putative sterol biosynthesis gene identified through previous BLASTp searches. N/A denotes organisms without listed sterol biosynthesis gene. --- denotes no homolog found with set e-value cutoffs.

**Table S2.** Mutations present in serial passaged *Calothrix* sp. NIES-4105.

| Evidence | Mutation | Annotation | Gene | Description |
|---|---|---|---|---|
| RA | Δ1 bp | intergenic (+33/-65) | *118839 → / → 118838* | hypothetical protein/hypothetical protein |
| RA | (C)11→10 | intergenic (-194/+40) | *114372← / ← 114371* | hypothetical protein/hypothetical protein |
| RA | C→G | intergenic (-346/+392) | *114310← / ← 114309* | 3-octaprenyl-4-hydroxybenzoate carboxy-lyase/3'-5' exoribonuclease, VacB and RNase II |
| RA | 2 bp→GA | intergenic (-385/+352) | *114310← / ← 114309* | 3-octaprenyl-4-hydroxybenzoate carboxy-lyase/3'-5' exoribonuclease, VacB and RNase II |
| RA | +G | coding (2515/2589 nt) | *114218 ←* | TPR repeat-containing protein |
| RA | 4 bp→AGAA | coding (2511-2514/2589 nt) | *114218←* | TPR repeat-containing protein |
| RA | G→C | L709L (CTC→CTG) | *113884 ←* | TPR repeat-containing protein |
| JC | +34 bp | coding (2106/2169 nt) | *113884 ←* | TPR repeat-containing protein |
| RA | Δ1 bp | coding (991/1116 nt) | *113884 ←* | hypothetical protein |
| JC | 13 bp→130 bp | coding (579-591/951 nt) | *113087←* | hypothetical protein |
| RA | T→G | intergenic (+303/+143) | *112194 → / ← 112193* | hypothetical protein/hypothetical protein |
| RA | T→C | intergenic (+323/-312) | *111735 → / → 11734* | 50S ribosomal protein L25/hypothetical protein |
| RA | T→G | intergenic (+328/-307) | *111735 → / → 11734* | 50S ribosomal protein L25/hypothetical protein |
| RA | A→T | intergenic (-445/-271) | *119725 ← / → 119724* | hypothetical protein/hypothetical protein |

RA indicates read alignment evidence for mutation. JC indicates new junction evidence for mutation. Mutation column describes specific mutation. In intergenic mutations, the distance from the start (+) or stop (-) of nearest genes are listed. In coding mutations, the location of mutation in gene as well as original nucleotide length is listed. IMG locus tags and annotations for each gene or nearest genes are listed.

**Table S3.** Bacterial strains used in this study.

| Strain | Genotype/Description | Source |
|---|---|---|
| *Enhygromyxa salina* DSM 15201 | Wild type DSM 15201 | DSMZ |
| *Calothrix* sp. NIES-4105 | Wild type NIES-4105 | NIES |
| *Escherichia coli* DH10B | Strain used for constructing plasmids, heterologous expression, and culturing *E. salina.*<br><br>$F^-$endA1 recA1 galE15 galK16 nupG rpsL ΔlacX74 Φ80lacZΔM15 araD139 Δ(ara,leu)7697 mcrA Δ(mrr-hsdRMSmcrBC)$\lambda^-$ | Invitrogen |

**Table S4.** Oligonucleotides used in this study.

| Oligonucleotide | Sequence | Notes |
|---|---|---|
| AL1 | CAATTTCACACAGGAGGCAAGCATATGAGCC GATCGATCAGAAAC | SLIC-pSRK-NdeI-MCAT *sdmA* F |
| AL2 | CGCGCTTGGCGTAATCATGGTCATCATGCCG GGTCTGCC | SLIC-pSRK-NdeI-MCAT *sdmA* R |
| AL3 | CAATTTCACACAGGAGGCAAGCATATGACCA CACTGGTCACCGGC | SLIC-pSRK-NdeI-MCAT *sdmB* F |
| AL4 | GCGCTTGGCGTAATCATGGTCATCAGATCAT CCCCCTCTCCCT | SLIC-pSRK-NdeI-MCAT *sdmB* R |
| AL5 | AATGCAGCTGGCACGACAGG | pSRK seq F |
| AL7 | CCAGGGTTTTCCCAGTCAC | pSRK seq R |
| AL22 | CCGCCAGGCAAATTCTGTTT | pBAD seq R |
| AL23 | CGTCACACTTTGCTATGCCA | pBAD seq F |
| AL34 | TTCTTTCCGAAGGCGTCGC | ESA *sdmB* seq |
| AL35 | TTGCCCCAAAAGGTGACCTCG | ESA *sdmA* seq |
| AL99 | TTGGGCTAGCAGGAGGAATTCACATGTCTAC CAAAGTTCGCATCCCC | SLIC-pBAD-NcoI-ESA *sdmA* F |
| AL100 | GACTCTAGAGGATCCCCGGGTAC TCACGACCCGCTGGGC | SLIC-pBAD-NcoI-ESA *sdmA* R |
| AL95 | TTGGGCTAGCAGGAGGAATTCACATGAGTGA AGCCGAACCCACAG | SLIC-pBAD-NcoI-ESA *sdmB* F |
| AL96 | GACTCTAGAGGATCCCCGGGTACTTACTCGG CGGCTTCGGT | SLIC-pBAD-NcoI-ESA *sdmB* R |
| AL111 | CTGTGGGTTCGGCTTCACTCATTTACGACCC GCTGGGCA | SLIC-pBAD-NcoI-ESA *sdmA*-overlap w *sdmB*-R |
| AL112 | TGCCCAGCGGGTCGTAAATGAGTGAAGCCG AACCCACAG | SLIC-pBAD-NcoI-ESA *sdmB*-overlap w *sdmA*-F |
| AL142 | GCCCGGGGGATCCACTAGTTTCAGTGTCGG CTCGAGAT | SLIC-pBAD-XbaI-ESA *sdmC* R |
| AL143 | CCACCGCGGTGGCGGCCGCTATGGCTGACC CAGCGTAT | SLIC-pBAD-XbaI-ESA *sdmC* F |
| AL161 | TTCGCGCGCTGTTTCAGCC | ESA sdmC seq |
| AL198 | TTGGGCTAGCAGGAGGAATTCACATGAAAGA CATAGCTATAAAAGGCG | SLIC-pBAD-NcoI-CALO *sdmA* F |
| AL199 | GACTCTAGAGGATCCCCGGGTACCTAGTTTG GGGCGCTCG | SLIC-pBAD-NcoI-CALO *sdmB* R |
| AL200 | CTAAGTTTTCTGACTGTTGA | CALO sdmAB Seq |
| AL258 | GACTCTAGAGGATCCCCGGGTACTCAACAGT CAGAAAACTTAGCCTCA | SLIC-pBAD-NcoI-CALO *sdmA* R |
| AL259 | AGCTCCCGTTACCAGAATTGTCATTCAACAGT CAGAAAACTTAGCCTCA | Overlap PCR CALO *sdmA* R |
| AL260 | TTGGGCTAGCAGGAGGAATTCACATGACAAT TCTGGTAACGGGAG | SLIC-pBAD-NcoI-CALO *sdmB* F |
| AL261 | GCTAAGTTTTCTGACTGTTGAATGACAATTCT GGTAACGGGAG | Overlap PCR CALO *sdmB* F |

| | | |
|---|---|---|
| AL264 | CAATTTCACACAGGAGGCAAGCATATGTCTG AACATTTAAACACCAAAC | SLIC-pBAD-NcoI-CALO *osc*-F |
| AL265 | CGCGCTTGGCGTAATCATGGTCATCATCAAA CCATTCGTTTCAACCG | SLIC-pBAD-NcoI-CALO *osc*-R |

F indicates forward primer, R indicates reverse primer, seq indicates sequencing primer, MCAT indicates *Methylococcus capsulatus*, ESA indicates *Enhygromyxa salina*, and CALO indicates *Calothrix* sp. NIES_4105.

**Table S5.** Plasmids used in this study.

| Plasmid | Description | Reference |
|---|---|---|
| pTrc-*sqs*-synRBS-*osc*-synRBS-*smo* (pABB501) | MEALZ_3096-MEALZ_0768-MEALZ_0767 (Squalene synthase-Oxidosqualene cyclase-Squalene epoxidase) optimized expression plasmid (altered osc and smo RBSs). | Lee et al, 2018 |
| pSRKGm-lacUV5-rbs5 (pABB492) | pBBR1 ori, lacUV5 promoter, Gmr | Banta et al, 2017 |
| pBAD1031K (pABB466) | pRV1031 ori, pBAD promoter, Kanr, | Chakravartty and Cronan, 2015 |
| pSRK_MCAT_*sdmA* (pAL7004) | H156DRAFT_2756 (*sdmA*) expression plasmid<br><br>H156DRAFT_2756 was amplified by PCR with primers AL1 and AL2. The fragment was assembled by SLIC into the NdeI site of pABB492. Sequence was confirmed with oligos AL5 and AL7. | (*) |
| pSRK_MCAT_*sdmB* (pAL7003) | H156DRAFT_2755 (*sdmB*) expression plasmid<br><br>H156DRAFT_2755 was amplified by PCR with primers AL3 and AL4. The fragment was assembled by SLIC into the NdeI site of pABB492. Sequence was confirmed with oligos AL5 and AL7. | (*) |
| pBAD1031K_ESA_*sdmA* (pAL7134) | Ga0097779_103152 (*sdmA*) expression plasmid<br><br>Ga0097779_103152 was amplified by PCR with primers AL99 and AL100. The fragment was assembled by SLIC into the NcoI site of pABB466. Sequence was confirmed with oligos AL22 and AL23. | (*) |
| pBAD1031K_ESA_*sdmB* (pAL7169) | Ga0097779_103151 (*sdmB*) expression plasmid<br><br>Ga0097779_103151 was amplified by PCR with primers AL95 and AL96. The fragment was assembled by SLIC into the NcoI site of pABB466. Sequence was confirmed with oligos AL22 and AL23. | (*) |
| pBAD1031K_ESA_*sdmC* (pAL7180) | Ga0097779_109097 (*sdmC*) expression plasmid<br><br>Ga0097779_103152 was amplified by PCR with primers AL142 and AL143. The fragment was assembled by SLIC into the | (*) |

| | | |
|---|---|---|
| | XbaI site of pABB466. Sequence was confirmed with oligos AL22 and AL23. | |
| pBAD1031K_ESA_*sdmAB* (pAL7141) | Ga0097779_103152 (*sdmA*) and Ga0097779_103151 (*sdmB*) co-expression plasmid<br><br>Ga0097779_103152 and Ga0097779_103151 were amplified by PCR with primers AL99, AL111 and AL96 and AL112, respectively. Fragments were annealed using overlap extension PCR and assembled by SLIC into the NcoI site of pABB466. Sequence was confirmed with oligos AL22, AL23, AL34 and AL35. | (*) |
| pBAD1031K_ESA_*sdmAC* (pAL7181) | Ga0097779_103152 (sdmA) and Ga0097779_109097(sdmC) co-expression plasmid<br><br>Ga0097779_109097 was amplified using oligos AL142 and AL143. The fragment was assembled by SLIC into the XbaI site of pAL7134. Sequence was confirmed with oligos AL22 and AL161. | (*) |
| pBAD1031K_ESA_*sdmAB-sdmC* (pAL7170) | Ga0097779_103152 (sdmA), Ga0097779_103151 (sdmB), and Ga0097779_109097(sdmC) co-expression plasmid<br><br>Ga0097779_109097 was amplified using oligos AL142 and AL143. The fragment was assembled by SLIC into the XbaI site of pAL7141. Sequence was confirmed with oligos AL22, AL23 and AL161. | (*) |
| pBAD1031K_CALO_*sdmA* (pAL7298) | Ga0263810_115380 (sdmA) expression plasmid<br><br>Ga0263810_115380 was amplified by PCR with primers AL198 and AL258. The fragment was assembled by SLIC into the NcoI site of pABB466. Sequence was confirmed with oligos AL22 and AL23. | (*) |
| pBAD1031K_CALO_*sdmB* (pAL7296) | Ga0263810_115382 (*sdmB*) expression plasmid<br><br>Ga0263810_115382 was amplified by PCR with primers AL199 and AL260. The fragment was assembled by SLIC into the NcoI site of pABB466. Sequence was confirmed with oligos AL22 and AL23. | (*) |

| | | |
|---|---|---|
| pBAD1031K_CALO_*sdmAB* (pAL7297) | Ga0263810_115380 (*sdmA*) and Ga0263810_115382 (*sdmB*) co-expression plasmid<br><br>Ga0263810_115380 and Ga0263810_115382 were amplified by PCR with primers AL198, AL259 and AL199 and AL261, respectively. Fragments were annealed using overlap extension PCR and assembled by SLIC into the NcoI site of pABB466. Sequence was confirmed with oligos AL22, AL23 and AL200. | (*) |
| pSRK_CALO_*osc* (pAL7309) | Ga0263810_115390 (*osc*) expression plasmid<br><br>Ga0263810_115390 was amplified by PCR with primers AL264 and AL265. The fragment was assembled by SLIC into the NdeI site of pABB492. Sequence was confirmed with oligos AL5 and AL7. | (*) |

(*) indicates this study, RBS indicates ribosome binding site, MCAT indicates *Methylococcus capsulatus*, ESA indicates *Enhygromyxa salina*, and CALO indicates *Calothrix* sp. NIES_4105.

**Table S6.** Heterologous expression strains used in this study.

| Expression Strain | Plasmids |
|---|---|
| PVW 7011 | pJBEI2997 (pABB302), pTrc-*sqs*-synRBS-*osc*-synRBS-*smo* (pABB501), pSRKGm-lacUV5-rbs5 (pABB492), pBAD1031K (pABB466) |
| PVW 7130 | pJBEI2997 (pABB302), pTrc-*sqs*-synRBS-*osc*-synRBS-*smo* (pABB501), pBAD1031K_ESA_*sdmA* (pAL7134), pSRK_MCAT_*sdmB* (pAL7003) |
| PVW 7131 | pJBEI2997 (pABB302), pTrc-*sqs*-synRBS-*osc*-synRBS-*smo* (pABB501), +pBAD ESA 1017 + pSRK_MCAT_*sdmA* (pAL7004) |
| PVW 7305 | pJBEI2997 (pABB302), pTrc-*sqs*-synRBS-*osc*-synRBS-*smo* (pABB501), pSRKGm-lacUV5-rbs5 (pABB492), pBAD1031K_ESA_*sdmA* (pAL7134) |
| PVW 7119 | pJBEI2997 (pABB302), pTrc-*sqs*-synRBS-*osc*-synRBS-*smo* (pABB501), pSRKGm-lacUV5-rbs5 (pABB492), pBAD1031K_ESA_*sdmB* (pAL7169) |
| PVW 7228 | pJBEI2997 (pABB302), pTrc-*sqs*-synRBS-*osc*-synRBS-*smo* (pABB501), pSRKGm-lacUV5-rbs5 (pABB492), pBAD1031K_ESA_*sdmC* (pAL7180) |
| PVW 7143 | pJBEI2997 (pABB302), pTrc-*sqs*-synRBS-*osc*-synRBS-*smo* (pABB501), pSRKGm-lacUV5-rbs5 (pABB492), pBAD1031K_ESA_*sdmAB* (pAL7141) |
| PVW 7303 | pJBEI2997 (pABB302), pTrc-*sqs*-synRBS-*osc*-synRBS-*smo* (pABB501), pSRKGm-lacUV5-rbs5 (pABB492), pBAD1031K_ESA_*sdmAC* (pAL7181) |
| PVW 7304 | pJBEI2997 (pABB302), pTrc-*sqs*-synRBS-*osc*-synRBS-*smo* (pABB501), pSRKGm-lacUV5-rbs5 (pABB492), pBAD1031K_ESA_*sdmAB-sdmC* (pAL7170) |
| PVW 7301 | pJBEI2997 (pABB302), pTrc-*sqs*-synRBS-*osc*-synRBS-*smo* (pABB501), pSRKGm-lacUV5-rbs5 (pABB492), pBAD1031K_CALO_*sdmA* (pAL7298) |
| PVW 7300 | pJBEI2997 (pABB302), pTrc-*sqs*-synRBS-*osc*-synRBS-*smo* (pABB501), pSRKGm-lacUV5-rbs5 (pABB492), pBAD1031K_CALO_*sdmB* (pAL7296) |
| PVW 7299 | pJBEI2997 (pABB302), pTrc-*sqs*-synRBS-*osc*-synRBS-*smo* (pABB501), pSRKGm-lacUV5-rbs5 (pABB492), pBAD1031K_CALO_*sdmAB* (pAL7297) |
| PVW 7310 | pJBEI2997 (pABB302), pTrc-*sqs*-synRBS-*osc*-synRBS-*smo* (pABB501), pSRK_CALO_*osc* (pAL7309), pBAD1031K (ABB466) |

All expression strains are *E. coli* DH10B with pJBEI2997 (pABB302, CmR), pTrc (pABB278 or derivatives, AmpR),pSRK (pABB492 or derivatives, Gmr) (where indicated), and pBAD1031K (pABB466 or derivatives, KanR) (where indicated).

**Dataset S1 (separate file).** Sterol biosynthesis pathway homologs in bacteria. We conducted a BLASTp search of bacteria in the genomic databases on the JGI IMG portal for oxidosqualene cyclase (1 x e $^{-50,}$ 30%ID). Of this subset of bacteria, we conducted further BLASTp searches using the putative cholesterol biosynthesis genes from *E. salina* and *Calothrix*. Cultured bacteria are bolded. IMG locus tags are listed for and highlighted green for organisms which harbor homologs. SMO, squalene monooxygenase; OSC, oxidosqualene cyclase; CYP51, C-14 demethylase; SdmA, sterol demethylase A; SdmB, Sterol demethylase B; SdmC, sterol demethylase C.

7. , Flux analysis of cholesterol biosynthesis in vivo reveals multiple tissue and cell-type specific pathways - PMC (August 30, 2022).
